# MLL2 facilitates long-range gene regulation through LINE1 elements

**DOI:** 10.1101/2025.08.10.669526

**Authors:** Lara Zorro Shahidian, Lucio Di Filippo, Sarah Malika Robert, Alvaro Rada-Iglesias

**Author notes:** Equal contribution (Lara Zorro Shahidian, Lucio Di Filippo).

## Abstract

Transcriptional regulation is tightly linked to chromatin organization, with H3K4me3 commonly marking both active and bivalent promoters. In embryonic stem cells (ESC), MLL2 is essential for H3K4me3 deposition at bivalent promoters, which has been proposed to facilitate the induction of major developmental genes during pluripotent cell differentiation. However, prior studies point to a functional discrepancy between the loss of H3K4me3 at bivalent promoters and the largely unaltered transcription of major developmental genes in *Mll2*^-/-^ cells. In this study, we investigated MLL2-dependent gene regulation in mouse ESC and during their differentiation. Contrary to the prevailing view, we show that MLL2’s primary role is not to oppose Polycomb-mediated repression at the bivalent promoters of developmental genes. Instead, we identify a previously unrecognized regulatory function for MLL2 at the CG-rich 5’ untranslated regions (5’UTR) of evolutionarily young LINE-1 (L1) transposable elements (TE). We found that MLL2 binds to the 5’UTR of L1 elements and is critical for maintaining their active state (H3K4me3 and H3K27ac), while preventing the accumulation of repressive H3K9me3. Using both global genomic approaches (i.e. RNA-seq, ChIP-seq and Micro-C) as well as targeted L1 deletions, we demonstrate that these MLL2-bound L1 elements act as enhancers, modulating the expression of neighboring genes in ESC and, more prominently, during differentiation. Together, our findings illuminate novel aspects of MLL2 regulatory function during early developmental transitions and highlight the emerging role of TE as key components of long-range gene expression control.

## Introduction

Transcriptional regulation largely depends on proper chromatin organization^1^. Histone modifiers, together with DNA methyltransferases, demethylases, and chromatin remodeling complexes, control chromatin accessibility, ensuring the correct environment is established to accommodate chromatin function at a given locus ^2^. The regulatory function of histone modifiers is closely linked to their enzymatic capacity to establish histone posttranslational modifications (PTM)^3^. These modifications can directly influence chromatin structure and accessibility and/or serve as recruitment platforms for effector complexes that control chromatin properties and gene expression (as reviewed in ^4^).

Among these modifications, methylation of lysine 4 on histone H3 (H3K4) is an evolutionarily conserved mark associated with active transcription^5^. It exists in three distinct states — mono-, di-, and trimethylation (H3K4me1/2/3) — which are differentially distributed across various types of regulatory elements. H3K4me1 is typically associated with enhancers: active enhancers are marked by both H3K4me1 and H3K27ac, while primed enhancers are enriched in H3K4me1 alone^6,7^. The importance of H3K4me1 for enhancer function remains questionable and likely depends on the cellular context^8^. H3K4me3 is commonly enriched at active promoters^7,9^, where recent evidence suggests that it directly facilitates RNA polymerase II (polII) pause-release and elongation^10^. In addition to marking active promoters, H3K4me3 is also enriched at bivalent promoters^11^, which are characterized by the co-occurrence of active (H3K4me3) and repressive (H3K27me3) marks on the same nucleosomes^11^. This seemingly contradictory chromatin signature was first described in embryonic stem cells (ESC), where it preferentially marks promoters of inactive genes with major developmental functions^12–14^. Based on these observations, it has been proposed that chromatin bivalency confers developmental genes a poised transcriptional state that prepares them for either full activation or repression upon differentiation and lineage commitment^11,15^. However, there is growing evidence that challenges several aspects of this model. First, bivalent promoters are also found in differentiated cell types such as CD4+ T cells and neurons^16–18^, suggesting their role might extend beyond developmental transitions. Second, and more importantly, although many bivalent promoters are bound by paused RNA polII phosphorylated at Ser5^19,20^, bivalency does not appear to confer faster gene induction dynamics but might rather protect promoters from *de novo* DNA methylation^21^. Together these findings suggest that bivalent chromatin may serve broader functions, contributing to gene regulatory flexibility and adaptation across diverse cellular contexts ^22,23^.

The majority of bivalent promoters contain CpG islands (CGI), that enable the recruitment of the histone methyltransferase complexes responsible for the establishment of H3K27me3 and H3K4me3^24,25^. H3K27me3 is catalyzed by Polycomb Repressive Complex 2 (PRC2), which together with the Polycomb Repressive Complex 1 (PRC1), maintains bivalent promoters in a transcriptional inactive state^26^. H3K4 methylation, in contrast, can be established by six different mammalian complexes, including MLL1-4 (KMT2A-D) COMPASS-like complexes and SET1A/B (KMT2F/G) COMPASS complexes (as reviewed by ^27^). In ESC, SET1A/B complexes preferentially bind CGI-rich active promoters, establishing and maintaining H3K4me3 at these sites^28^. In contrast, the MLL2 COMPASS-like complex binds most CGI-rich promoters in ESC, regardless of their transcriptional status, yet it is only required for H3K4me3 at bivalent promoters^28–31^. In ESC, MLL1 plays a redundant role that only gets manifested in the absence of MLL2^31^, while in other cell types, both MLL1 and MLL2 complexes contribute to H3K4me3 deposition at bivalent regions^32^. MLL1 and MLL2 are homologous to the *Drosophila* protein Trithorax (Trx) and share a conserved domain architecture. Both proteins contain: (i) a CXXC domain (which specifically recognizes unmethylated CpG-rich sequences and, thus, preferentially targets MLL1/2 complexes to CGI^29^), an AT hook (involved in DNA binding and protein interactions^33^), four N-terminal plant homeotic domains (PHD, that mediate binding to methylated H3^34^), and a C-terminal SET domain (which catalyzes H3K4 methylation^35^). MLL1 and MLL2 form multi-protein COMPASS-like complexes that include the core WRAD (WDR5, RBBP5, ASH2L, and DPY30) module, as well as HCF1, MENIN, and LEDGF. The WRAD core module regulates enzymatic activity, stabilizes the complex and facilitates its recruitment to chromatin^36^. HCF1 further enhances stability^37^, while MENIN mediates locus-specific targetting^38^. Additionally, the MLL1 COMPASS-like complex includes the co-activator LEDGF, that is absent in other complexes^39^.

MLL2 plays a central role in depositing H3K4me3 at bivalent promoters in pluripotent cells both *in vitro* and *in vivo* ^29–31,40^. According to the proposed role of promoter bivalency in gene poising ^11,15,41^, the loss of MLL2 function should impair the induction of major developmental genes during pluripotent cell differentiation. In agreement with this prediction, *Mll2*^-/-^ mice show early developmental and growth defects resulting in embryonic lethality before E11.5^42^. Nonetheless, transcriptional profiling of both undifferentiated and differentiated *Mll2*^-/-^ ESC revealed important discrepancies between gene expression changes and the loss of H3K4me3 at bivalent promoters^30,31,43,44^. Firstly, the overall number of downregulated genes in *Mll2^-/-^* ESC is considerably smaller than the number of bivalent genes that lose H3K4me3 in the absence of MLL2^30,31,43,44^. Consequently, only a relatively small fraction of all bivalent genes (<10%) becomes downregulated in *Mll2^-/-^*ESC^30,44^. Furthermore, less than half of all downregulated genes in *Mll2*^-/-^ ESC can be classified as bivalent^30^. Similarly, a minority (<25%) of bivalent genes that are normally induced during ESC differentiation fail to do so in the absence of MLL2^30^. It has been suggested that the reduced expression of bivalent genes in the absence of MLL2 could be the result of the antagonistic regulatory functions of MLL2 and Polycomb complexes in the context of bivalent chromatin^30,45^. In *Drosophila*, trithorax group (TrxG) and polycomb group (PcG) proteins co-regulate homeotic genes (Hox) with opposing roles: TrxG activate their expression and PcG represses them (as reviewed by ^46^). In mammals, both complexes are recruited to hypomethylated CGI and it has been suggested they compete for these sites^47^. In agreement with this model, the loss of MLL2 in ESC leads to increased PRC2 and PRC1 binding and elevated H3K27me3 levels at bivalent promoters^30^. Nevertheless, the changes in H3K27me3 are generally moderate, particularly at the promoters of major developmental genes, such as the Hox genes^30,31,44,45^. The partial overlap between bivalent genes and genes whose expression is controlled by MLL2, together with the moderate increase in H3K27me3 levels that *Mll2*^-/-^ ESC show at bivalent promoters, suggest that the regulatory function of MLL2 in ESC and/or during their differentiation might involve additional mechanisms beyond antagonizing Polycomb-mediated repression at promoter regions. Several factors may account for this complexity:

(i) Previous work indicates that MLL2 and MLL1 share partially redundant functions in controlling H3K4me3 levels at bivalent promoters both in ESC, as well as upon their differentiation^31^. Therefore, in *Mll2*^-/-^ cells, the presence of MLL1 might protect, at least partially, bivalent promoters from the repressive effect of PcG complexes. However, neither gene expression nor H3K27me3 levels have been investigated in *Mll1*^-/-^; *Mll2*^-/-^ cells^31^.
(ii) Poised enhancers (PE) represent a unique set of distal regulatory elements that control the expression of major developmental genes both *in vivo* and *in vitro*^48,49^. Before becoming active and gaining H3K27ac in differentiating cells, PE display a characteristic bivalent chromatin signature in pluripotent cells that includes marking with H3K4me1/2 and H3K27me3 and binding of co-activators (e.g. p300) and PcG complexes^50^. Furthermore, PE display strong physical contacts with their target genes both before and after they become active upon pluripotent cell differentiation^49^. We recently showed that almost 80% of PE are associated with CGI, which, by acting as tethering elements that increase the physical and functional communication between the PE and their target genes, confer these enhancers with their unique chromatin, topological and regulatory properties^51^. It is currently unclear which protein complexes are recruited by the PE-associated CGI to boost enhancer activity and target gene expression. Nevertheless, the preferential binding of MLL2 to CGI, as well as its importance in controlling H3K4 methylation at bivalent loci, suggest that MLL2 could be an important regulator of the activity and/or topological features of PE and, thus, of major developmental genes.
(iii) It was recently reported that, in mESC, MLL2 binds to the CG-rich 5’UTR of young LINE-1 (L1) retrotransposons, thereby controlling their H3K4me3 levels and promoting their expression^52^. L1 retrotransposons are among the most abundant classes of transposable elements (TE) in the mammalian genome^53^. TE have gained significant attention in recent years due to their involvement in evolution and disease^54,55^, as well as in the control of transcription and 3D chromatin organization^56,57^. L1 are 6-7 Kb long and comprise a 5’-unstranslated region (5’-UTR), three open reading frames (ORF0, ORF1, ORF2), and a polyadenylated (polyA) 3’-UTR^53,58,59^. In mice, young L1 subfamilies, with high CG content in their 5’-UTR, remain active in certain cellular contexts despite host mechanisms designed to suppress them^60^. Host cells counter L1 activity through defense strategies, including nucleic acid editing, RNA-induced silencing, and epigenetic repression preferentially involving DNA and H3K9 methylation^61–63^. However, in specific cellular contexts, such as early developmental stages, brain corticogenesis or certain cancer types, L1 elements evade these defenses and become transcriptionally active, acting as alternative promoters, enhancers or lncRNA that participate in the control of gene expression programs^64–67^. Therefore, by binding to and activating young L1 elements (e.g. L1Md_A, L1Md_Gf and L1Md_T) with either *cis* or *trans* regulatory functions, MLL2 could potentially regulate gene expression in ESC and/or during their differentiation.

Here, we identify a previously unknown regulatory mechanism through which MLL2 controls gene expression in both ESC and, especially, their differentiated progeny. We demonstrate that MLL2 binding to the CG-rich 5’UTR of specific L1 subfamilies is essential for the enhancer-like activity of these elements and, thus, for the proper expression of nearby MLL2 target genes. These findings help reconcile previously unexplained observations regarding the role of MLL2 during pluripotent cell differentiation and underscore the emerging role of transposable elements as active contributors to long-range gene regulation.

## Results

### Characterization of the chromatin and transcriptional changes associated with the conditional loss of MLL2 in mouse pluripotent cells

To understand the mechanisms governing MLL2-dependent gene regulation in pluripotent cells, we set off to characterize chromatin and gene expression changes specifically associated with the loss of MLL2 in mESC. Although MLL1 is lowly expressed in mESC and its loss does not lead to major gene expression changes^31^, it has been reported that, in the absence of MLL2, MLL1 can partially compensate for its loss^31^. Accordingly, we investigated MLL2 function, both in the absence and in the presence of MLL1. Starting from a previously described Cre-inducible *Mll2*-KO mESC line^43^ (Fig. S1A-B), we generated constitutive *Mll1*-KO cells via CRISPR editing (Fig. S1C). This approach enabled us to compare chromatin and transcriptional features of WT, single *Mll1*-KO, single *Mll2*-KO, and double *Mll1*-KO/*Mll2*-KO (double-KO) mESC (Fig. 1A). To ensure that the observed changes were the direct effects of MLL2 loss, rather than the result of cellular adaptation, 4-hydroxytamoxifen (4-OHT) treatment was always performed 48 hours before each experiment.

**Figure 1.**
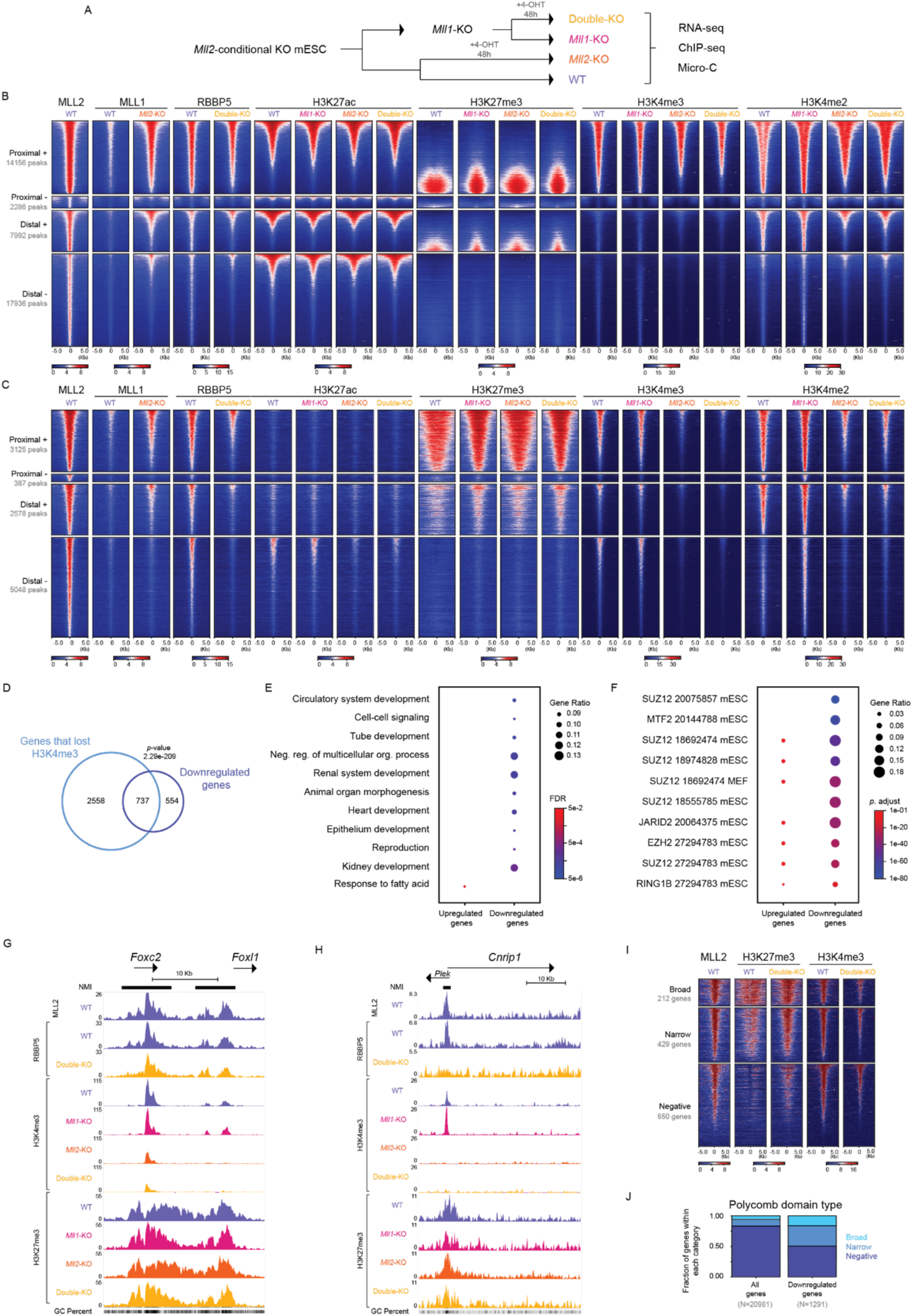
Chromatin and transcriptional changes upon MLL2 loss in mESC. (A) Schematic representation of the experimental workflow used to investigate the regulatory function of MLL2 in ESC. Starting from a Cre-inducible Mll2-KO mESC line, Mll1 was constitutively knocked out using CRISPR-Cas9 genome editing. Treatment with 4-OHT (final concentration of 800 nM, 48 hours) in WT and Mll1-KO cells resulted in the generation of Mll2-KO and double-KO mESC, respectively. All experiments were performed 48 hours after the 4-OHT treatment. (B) Heatmap plots showing the ChIP-seq signals for the indicated proteins and histone marks around all the MLL2 peaks identified in WT mESC. MLL2 peaks were categorized as: proximal +/- and distal +/-. Peaks were considered proximal if located within 5 Kb of a TSS; otherwise, they were classified as distal. Peaks were further designated as positive (+) or negative (-) based on proximity (within 1 Kb) to CpG islands (CGI). Within each category, the MLL2 peaks were ranked according to H3K27ac signals in WT ESC. (C) Heatmap plots showing the ChIP-seq signals for the indicated proteins and histone marks around those MLL2 peaks that lost H3K4me3 in the double-KO mESC. The MLL2 peaks were divided into the four categories described in (B) and, within each category, the peaks are ranked by the H3K4me3 signals in WT cells. (D) Overlap between genes associated with MLL2 proximal peaks losing H3K4me3 in double-KO mESC (i.e. genes that lost H3K4me3) and genes downregulated in double-KO mESC. The p-value was calculated using a hypergeometric test. (E) Gene ontology (GO) enrichment analysis for the “biological process” category was performed for up-(left) and downregulated (right) genes in double-KO mESC using WebGestalt^CS^. Representative examples from the top 20 most enriched terms are shown. (F) Enrichment analysis for the “ChEA 2022 transcription factor targets” was performed using Enrichr^70^ for up-(left) and downregulated (down) genes in double-KO mESC. Representative examples from the top 20 enriched terms are shown. (G-H) Genome browser snapshots showing the ChIP-seq profiles of MLL2, RBBP5, H3K4me3 and H3K27me3 in WT, Mll1-KO, Mll2-KO and double-KO mESC around (G) Foxc2 and Foxl1, two genes with broad PcG domains that are not downregulated in double-KO mESC despite losing H3K4me3 and (H) Cnrip1, a gene downregulated in double-KO mESC. Non-Methylated Islands (NMI)^71^ are shown as black rectangles at the top of the panels, while “GC percent” tracks are shown at the bottom. (I) Heatmap plots showing MLL2, H3K4me3 and H3K27me3 ChIP-seq signals in WT and double-KO mESC around the TSS of all the downregulated genes in double-KO mESC. The TSS are sorted by the H3K4me3 signals in WT mESC. The downregulated genes were divided into three categories based on the length of their associated Polycomb domains (see Methods): broad (≥C Kb), narrow (0< length <C Kb) and negative (0 Kb). (J) Distribution of genes according to their associated Polycomb domains (broad (≥C Kb), narrow (0< length <C Kb) and negative (0 Kb)) is shown for either all mouse genes (left; n=20S81) or the genes downregulated in double-KO mESC (right; n=12S1).

ChIP-seq experiments were performed to investigate the binding profiles of MLL2, MLL1 and RBBP5 in WT mESC. Moreover, ChIP-seq experiments against MLL1 and RBBP5 were also performed in single *Mll2*-KO and double-KO mESC, respectively (Fig. 1B). Additionally, ChIP-seq experiments for a panel of histone PTMs (i.e. H3K4me1/2/3, H3K27ac, H3K27me3) were conducted across all four genotypes to characterize chromatin changes resulting from MLL1/2 loss (Fig. 1B, Fig. S2A). Given that MLL2 was previously reported to bind to the 5’UTR of some L1 elements^52^, the ChIP-seq data was mapped using a pipeline compatible with the analysis of transposable elements^68^ (see Methods). For MLL2, we identified a total of 42370 peaks in mESC (Supplementary Data 1), which were divided into four different groups attending to their proximity to transcription start sites (TSS; proximal or distal) and CGI (+ or -): (i) proximal +, (ii) proximal -; (iii) distal +; (iv) distal - (Fig. 1B, Fig. S2A). Consistent with previous reports, almost 40% of the MLL2 peaks (n=16442) were located near TSS, the majority of which (86%) were associated with CGI (i.e. proximal +)^29,31^. As previously described, *Mll2*-KO mESC showed a strong reduction in H3K4me3 (and to a lesser degree in H3K4me2) specifically at bivalent promoters enriched in H3K27me3, while H3K4me3 levels at active promoters, enriched in H3K27ac, remained largely unaffected (Fig. 1B). Despite the high sequence homology between MLL1 and MLL2, MLL1 showed low enrichment at MLL2-bound regions in WT mESC. Interestingly, in the absence of MLL2, MLL1 binding significantly increased at the majority of MLL2-bound regions (Fig. 1B), supporting a potential compensatory effect. However, H3K4me3 levels at MLL2-bound bivalent promoters were similarly reduced in *Mll2*-KO and double-KO mESC, while being unaffected in *Mll1*-KO mESC (Fig. 1B), thus confirming that the MLL2 COMPASS-like complex is largely responsible for H3K4me3 at bivalent promoters. Furthermore, the binding of RBBP5, a core component of COMPASS/COMPASS-like complexes, was specifically reduced in double-KO mESC at bivalent promoters losing H3K4me3 in the absence of MLL2, while remaining unchanged at active promoters (Fig. 1B). Overall, these observations are in agreement with previous reports and confirm that MLL2 is specifically required for the proper recruitment of COMPASS-like complexes and the deposition of H3K4me3 in the context of bivalent promoters.

To further characterize the chromatin changes caused by the loss of MLL2, we specifically identified those MLL2 peaks that significantly lost H3K4me3 in double-KO mESC compared to WT mESC (n=11138; Fig. 1C, Fig. S2B-C; Supplementary Data 1). In agreement with the preliminary observations described above, the regions losing H3K4me3 in double-KO mESC showed similar reductions in this histone PTM in *Mll2*-KO cells, while being unaffected in *Mll1*-KO cells. Furthermore, and as previously reported, the majority of proximal promoters losing H3K4me3 in double-KO cells (n=3512; 32%) were enriched in H3K27me3 in WT mESC, thus classifying them as bivalent promoters, and overlapped with CGI (89% proximal + (n=3125); 11% proximal - (n=387); Fig. 1C, Fig. S2B-C). Regarding the distal MLL2 peaks losing H3K4me3 in double-KO mESC (68%; n=7626), approximately 34% of them (n=2578) were located close to CGIs (i.e. distal +; Fig. 1C). Similarly to proximal *+* regions, distal + regions were enriched in H3K27me3, thus showing the typical genetic (i.e. CGI) and epigenetic (i.e. bivalent chromatin) features associated with PE^50^. Intriguingly, the largest fraction of MLL2 peaks associated with reduced H3K4me3 levels in double-KO mESC (n=5048; 45% of all MLL2 peaks and 66% of the distal MLL2 peaks) belonged to the distal - category. Interestingly, despite being clearly bound by MLL2 in WT mESC, distal - regions lacked H3K27me3 and exhibited low H3K27ac levels (Fig. 1C, Fig. S2C), suggesting they are not regulated by PcG complexes. Altogether, these analyses showed that MLL2 COMPASS-like complexes are required for proper H3K4me3 not only at bivalent promoters, as previously reported, but also at a large number of distal regions whose identity and potential functionality will be addressed in following sections.

Next, to assess the gene expression changes associated with MLL1/2 loss, we carried out RNA-seq experiments in WT and the three mutant mESC lines (Fig. 1A; Supplementary Data 2). Principal component analysis (PCA) revealed a clear separation of the samples according to their genotype, suggesting significant transcriptional differences between the different cell lines (Fig. S3A). As expected, *Mll1*-KO cells exhibited the lowest number of differentially expressed genes (DEG), while double-KO cells showed the highest (Fig. S3B). Consistent with MLL2 mainly acting as an activator, the majority of DEG in the double-KO mESC were downregulated relative to WT controls (i.e. 1365 downregulated *versus* 301 upregulated genes; Fig S3C). A comparison of the downregulated genes across the three mutant cell lines revealed that 75% of *Mll1*-KO and 69% of *Mll2*-KO downregulated genes overlapped with genes downregulated in double-KO ESC, respectively (Fig. S3D). Nevertheless, 771 additional genes were exclusively downregulated in the double-KO ESC, accounting for over 55% of all downregulated genes in these cells (Fig. S3D). These differential gene expression analyses suggest that, despite the negligible role of MLL1 in controlling H3K4me3 levels (Fig. 1B-C), MLL1 and MLL2 display partially overlapping gene regulatory functions in mESC. These results, together with previous reports^31^, further supported the use of double-KO cells in this study.

In pluripotent cells, MLL2 has been proposed to regulate developmental genes with bivalent promoters by antagonizing the repressive function of PcG complexes^30^. However, this model is somehow difficult to reconcile with the discrepancy between the relatively small number of genes downregulated in *Mll2*-KO ESC compared to the number of bivalent genes that lose H3K4me3 in the absence of MLL2, as reported in several studies^29–31,43^. Consistently, although we observed a significant overlap between the genes losing H3K4me3 at their promoters and the downregulated genes in double-KO mESC (p-value = 2.29e-209), only 22% of the genes losing H3K4me3 were actually downregulated and 57% of the downregulated genes exhibited H3K4me3 loss (Fig. 1D). This discrepancy was also reflected in the Gene Ontology (GO) terms that were significantly enriched in each gene group (Fig. S4A, Fig. 1E): while genes losing H3K4me3 were strongly enriched for developmental terms as expected for genes with bivalent promoters (Fig. S4A), the downregulated genes were more moderately associated with them (Fig. 1E). Nevertheless, a large proportion of downregulated genes in the double-KO ESC displayed hallmark features of bivalent chromatin, including strong enrichments for H3K4me3, H3K27me3 and several PcG subunits at their promoter regions (Fig. 1F-I) and low expression levels in WT ESC (Fig. S4B). To understand why these downregulated genes were not as strongly linked to developmental GO terms as the genes losing H3K4me3 at their promoter regions, we examined their chromatin features in more detail. Visual inspection of the generated ChIP-seq profiles, revealed that the promoter regions of non-DEG losing H3K4me3 in double-KO ESC frequently overlapped with broad H3K27me3 domains (Fig. 1G), a typical feature of major developmental genes (Fig. S4C)^72–74^. In contrast, the promoter regions of downregulated genes losing H3K4me3 in the double-KO ESC often overlapped with narrow and weaker H3K27me3 domains (Fig. 1H; Fig. S4D). To test whether the previous observations could be generalized, we categorized the genes downregulated in double-KO ESC according to the length of the H3K27me3 domains overlapping with their promoter regions into “broad” (H3K27me3 domain >6 Kb), “narrow” (0 Kb< H3K27me3 domain <6 Kb), and “negative” (H3K27me3 domain = 0; Fig. 1I and Fig. S4E; Supplementary Data 2). Interestingly, although the downregulated genes showed a moderate enrichment in the broad category in comparison to all genes (Fig 1J), only 16.4% of the downregulated genes were associated with a broad H3K27me3 domain (Fig. 1I-J). Furthermore, as many as 33% and 50% of the downregulated genes were classified as either “narrow” or “negative”, respectively. Altogether, these results challenge the prevailing view that MLL2 primarily regulates major developmental genes in pluripotent cells by antagonizing PcG-mediated repression.

### Antagonizing PcG-mediated repression is not the main regulatory role of MLL2 in mESC

In mammals, both MLL1/2 COMPASS-like and PcG complexes bind to unmethylated CGI^29,75,76^. It has been proposed that in the absence of MLL2, PcG binding increases at bivalent promoters, leading to transcriptional repression^30^. Our initial analyses, however, challenged this notion as: (i) MLL2-bound promoters that lost H3K4me3 in the absence of MLL2 (i.e. in *Mll2*-KO and double-KO ESC) showed rather mild increases in H3K27me3 (Fig. 1C, Fig S2C); (ii) only 23% of the genes whose promoters lost H3K4me3 in double-KO ESC, and that often displayed a bivalent chromatin state in WT cells, became downregulated in the absence of MLL2 (Fig. 1C-D); (iii) 50% of the downregulated genes in double-KO ESC are not enriched in H3K27me3 in WT ESC and, thus, are not bivalent (Fig. 1I-J). Importantly, PRC1, rather than PRC2, was previously shown to be primarily responsible for PcG-mediated repression in ESC^77,78^. To further test whether the regulatory function of MLL2 in ESC preferentially entails antagonizing PcG-mediated repression, we performed ChIP-seq experiments in WT and double-KO mESC against RING1B (Fig. 2A-D and Fig. S5A-B), the catalytic subunit of PRC1. When considering the MLL2-bound regions that lost H3K4me3 in double-KO mESC, we found that, in WT ESC, PRC1 binding profiles resembled those of H3K27me3 and, thus, PRC1 was preferentially enriched at proximal + and distal + regions (Fig. 2A,C,D and Fig. S5A). However, although the double-KO ESC displayed a moderate increase in H3K27me3 at those proximal + and distal + MLL2-bound regions^30^, RING1 binding was almost unchanged (Fig. 2A,C,D and Fig. S5A). Next, we similarly evaluated the RING1B binding profiles at the promoters of genes that were downregulated in double-KO ESC (Fig. 2B-C and Fig. S5B). Here again, although H3K27me3 levels increased in double-KO cells, particularly at “broad” and “narrow” promoters, RING1B binding remained unchanged, suggesting that PRC1 is not involved in the repression of MLL2 target genes.

**Figure 2.**
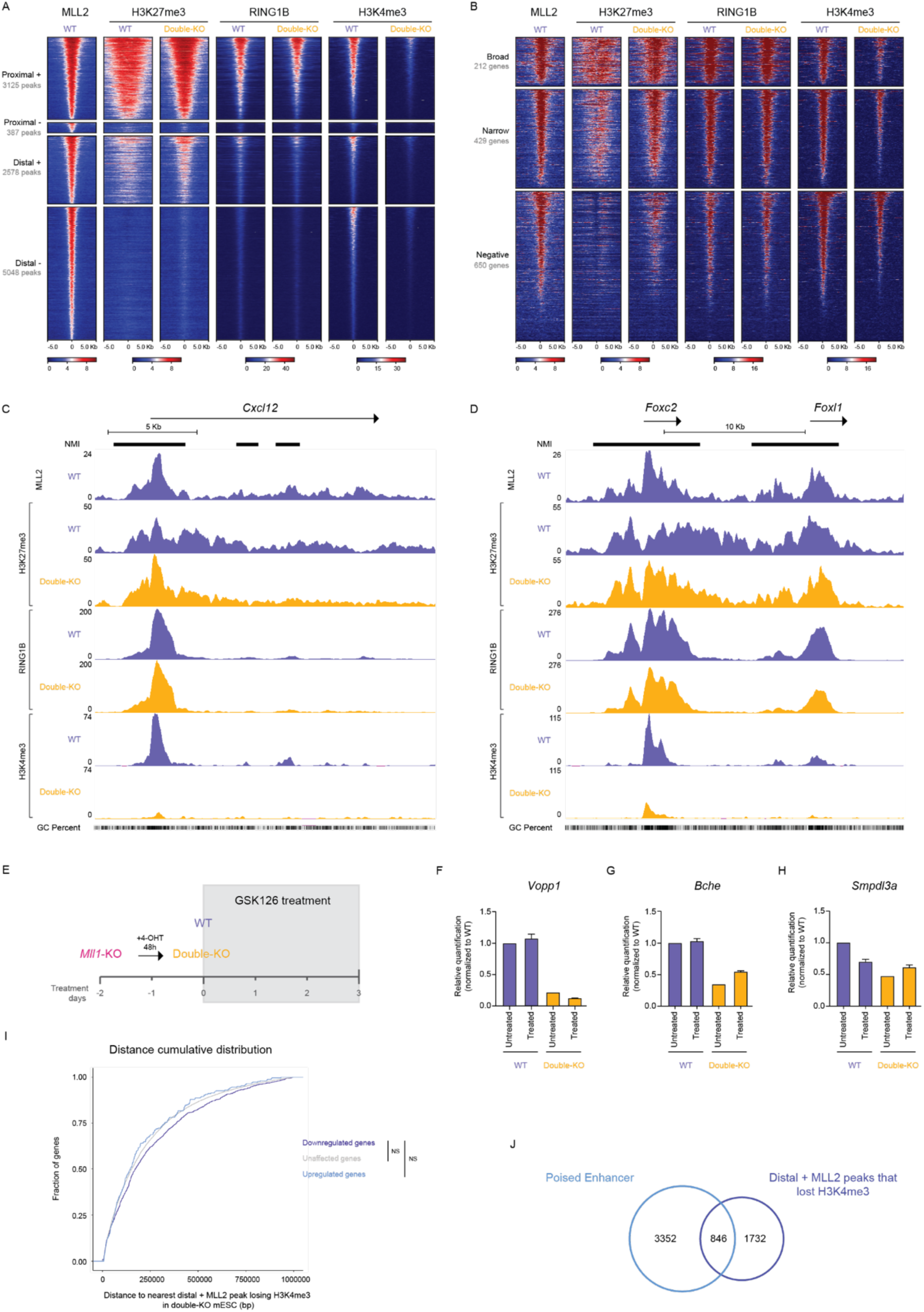
Increased Polycomb repression is not the main driver of the transcriptional changes associated with the loss of MLL2 in mESC. (A) Heatmap plots showing MLL2, H3K27me3, RING1B and H3K4me3 ChIP-seq signals in WT and double-KO mESC around MLL2 peaks that lost H3K4me3 in the double-KO mESC. MLL2 peaks are divided into four groups based on their distance to TSS (proximal or distal) and CGI (+ or -). Within each category MLL2 peaks are sorted by the H3K4me3 signals in WT mESC. (B) Heatmap plots showing MLL2, H3K27me3, RING1B and H3K4me3 ChIP-seq signals in WT and double-KO mESC around the TSS of all downregulated genes in double-KO mESC. The downregulated genes were divided into three categories based on the length of their associated Polycomb domains: broad (≥C Kb), narrow (0< length <C Kb) and negative (=0 Kb; see Methods). Within each category the TSS are sorted by the H3K4me3 signals in WT mESC. (C-D) Genome browser snapshots showing the ChIP-seq profiles of MLL2, H3K27me3, RING1B and H3K4me3 in WT and double-KO mESC around (C) Cxcl12, a gene downregulated in double-KO mESC and (H) Foxc2 and Foxl1, two genes with broad PcG domains that are not downregulated in double-KO mESC despite losing H3K4me3. NMI are shown as black rectangles at the top of the panels, while GC percent tracks are shown at the bottom. (E) Schematic representation of the GSK12C treatment workffow. Single Mll1-KO mESC were treated with 800 nM 4-OHT for 48 hours to generate double-KO mESC. Following the 4-OHT treatment, WT and double-KO mESC were treated with GSK12C (final concentration of 5 µM) for 72 hours. (F-H) The expression levels of (F) Vopp1, (G) Bche, and (H) Smpdl3a were measured by RT-qPCR in WT (purple) and double-KO (yellow) mESC with and without GSK12C treatment. Expression values are normalized with respect to untreated WT samples using two housekeeping genes (i.e. Hprt and Eef1a1) as loading controls. Two biological replicates were performed per condition. (I) Cumulative frequency distribution plot for the distances from the TSS of unaffected, upregulated and downregulated genes in double-KO mESC to the nearest distal + MLL2 peak losing H3K4me3 in double-KO mESC. The p-values were calculated using a Wilcoxon Rank-Sum test. NS: not significant. (J) Overlap between pluripotency-associated poised enhancers^48^ and distal + MLL2 peaks that lost H3K4me3 in double-KO mESC.

To further test whether the increase in H3K27me3 levels observed upon loss of MLL2 in the promoter regions of genes downregulated in double-KO ESC contributes to their repression, we treated WT and double-KO cells with GSK126, a well-characterized PRC2 inhibitor^79^ (Fig. 2E and Fig. S5C). GSK126 treatment effectively reduced H3K27me3 levels in both genotypes (Fig. S5D), confirming the efficacy of the inhibitor. We then analyzed the expression of three representative genes that were downregulated in double-KO ESC compared to WT ESC (Fig. 2F-H). If PRC2 activity was primarily responsible for the transcriptional repression of MLL2 target genes, we would expect GSK126 treatment to restore normal expression levels in double-KO ESC. RT-qPCR analyses confirmed the reduced expression of the three selected genes in untreated double-KO cells compared to untreated WT controls. However, the silencing of these three genes in the double-KO ESC was not restored by the GSK126 treatment. These findings further support that, although the promoter regions of genes downregulated in double-KO ESC display increased PRC2 binding and H3K27me3 deposition in the absence of MLL2, this is not the only or even the primary mechanism driving their repression.

In addition to proximal promoter regions, MLL2 also binds a large number of distal regions losing H3K4me3 in double-KO cells, including distal + regions resembling PE (Fig. 1C)^49,50^. We hypothesized that the regulatory activity of PE in ESC, although low, could be dependent, at least partly, on MLL2. This in turn could explain why lowly expressed genes in WT ESC get further repressed in double-KO ESC (Fig. S4B). If this hypothesis is correct, the distal + regions losing H3K4me3 in double-KO ESC should be located at shorter linear distances from genes downregulated in double-KO ESC than from unaffected or upregulated genes. However, a cumulative distribution analysis revealed no significant distance differences between the three gene groups (Fig. 2I). Furthermore, although MLL2-bound distal + regions losing H3K4me3 in double-KO ESC displayed genetic and epigenetic features typical of PE, only 30% of them actually overlapped with PE previously mapped in mouse pluripotent cells^48^ (Fig. 2J). Taken together, we conclude that MLL2-bound distal + regions with PE-like features are not responsible for the transcriptional changes observed in ESC upon MLL2 loss.

### MLL2 binds to the 5’UTR of specific LINE-1 subfamilies in ESC and controls their chromatin state

Having ruled out a major involvement of PcG-mediated repression and PE-like elements in the transcriptional changes associated with the loss of MLL2 in ESC, we set off to investigate alternative mechanisms through which MLL2 could regulate gene expression. Given that nearly half (45%) of all the MLL2 peaks losing H3K4me3 in double-KO cells belonged to the distal - category (Fig. 1C), we decided to investigate them in more depth. Interestingly, upon visual inspection of the regulatory domains of genes downregulated in double-KO ESC, we noticed that they often contained distal - MLL2 peaks overlapping the 5’UTR of L1 elements (Fig. 3A-B and Fig. S6A). Notably, these 5’UTR sequences were CG-rich, bound by RBBP5, enriched in H3K4me3 and H3K27ac and devoid of H3K27me3, indicating that the associated L1 elements were potentially active in ESC. Moreover, in double-KO cells, these distal - MLL2 peaks showed loss of RBBP5 binding, along with decreased levels of H3K4me3 and H3K27ac (Fig. 3A-B and Fig. S6A). To determine whether these initial observations could be somehow generalized, we calculated the distance between each distal - MLL2 peak losing H3K4me3 in double-KO ESC and the 5’UTR of the nearest L1 element (Fig. 3C). Distal - MLL2 peaks were significantly closer to L1 elements than the other groups of MLL2 peaks losing H3K4me3 in double-KO ESC, with as many as 62.5% of the distal - peaks located within 1 Kb of a L1’s 5’UTR (Fig. 3C; Supplementary Data 1). Furthermore, the MLL2-bound L1 elements were particularly enriched in young L1 subfamilies, such as L1Md_T, L1Md_A and L1Md_Gf (Fig. 3D; Supplementary Data 1), thus in agreement with previous reports^52^. Visualization of the GC-content and ChIP-seq datasets generated in our different ESC lines around the 5’end of the MLL2-bound L1 elements globally confirmed our initial observations at a few individual loci (Fig. 3E and Fig. S6B-C): (i) the 5’UTR of the MLL2-bound L1 elements displayed a high GC content that mirrored the MLL2 binding patterns, thus offering a molecular explanation as to how MLL2 can get recruited to these repetitive loci through its CXXC domain^80^; (ii) the investigated L1 elements represent bona-fide targets of the COMPASS-like complex, as the loss of MLL2 significantly reduced RBBP5 binding; (iii) the loss of MLL2 resulted in a strong reduction not only in H3K4me3 levels, but also in H3K27ac, indicating that the promoters of at least some of these MLL2-bound L1 elements are active in ESC; (iv) the 5’end of the MLL2-bound elements displayed low levels of H3K27me3 in WT and double-KO ESC, thus indicating that the loss of an active chromatin state at these loci in the absence of MLL2 is not mediated by PcG.

**Figure 3.**
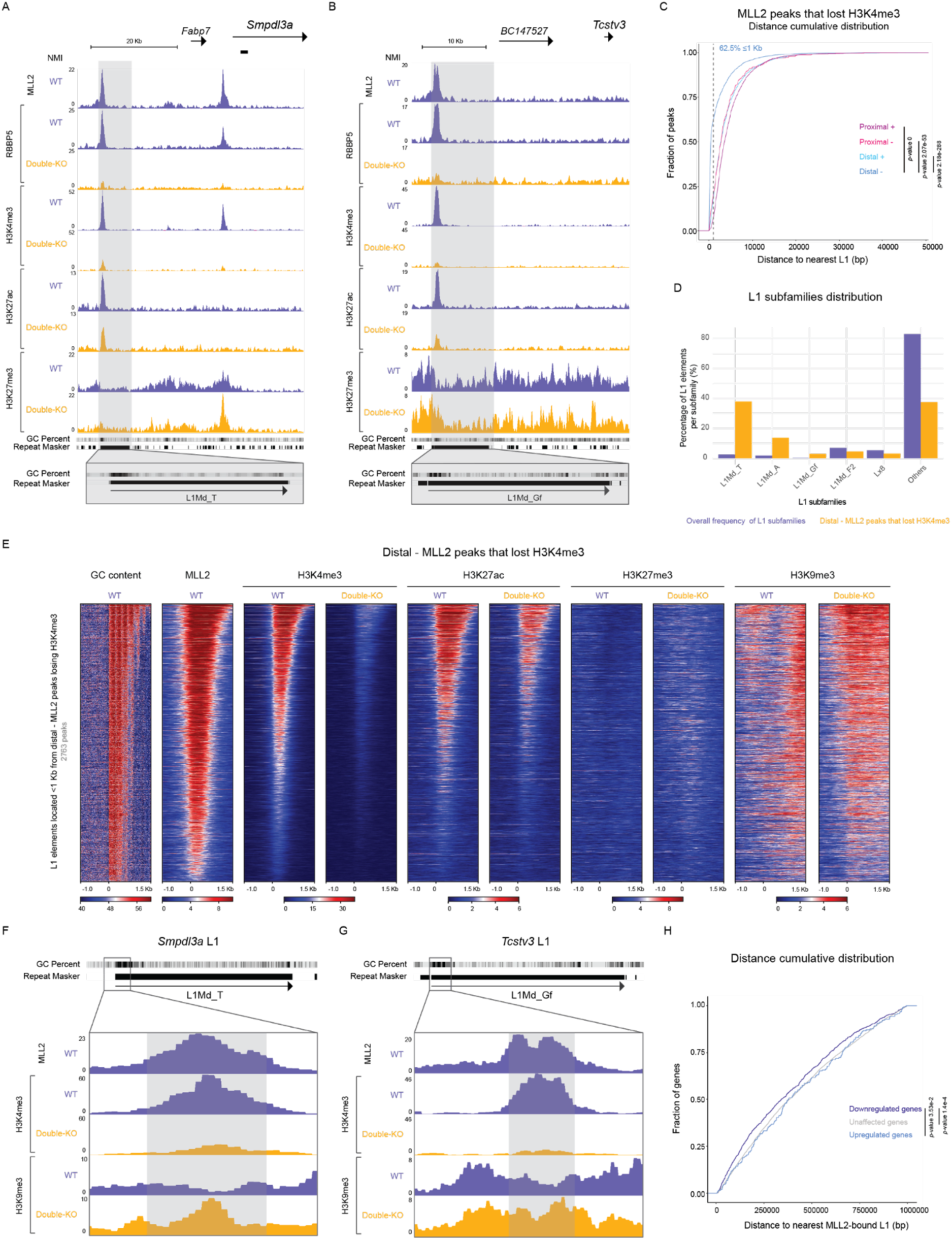
MLL2 binds to the GC-rich 5’UTRs of young L1 subfamilies and controls their chromatin features. (A-B) Genome browser snapshots showing the ChIP-seq profiles of MLL2, RBBP5, H3K4me3, H3K27ac and H3K27me3 in WT and double-KO mESC around (A) Smpdl3a and (B) Tcstv3, two genes that are downregulated in double-KO mESC. L1 elements bound by MLL2 are highlighted in grey. GC percent and Repeat Masker tracks are shown at the bottom. (C) Cumulative frequency distribution plot for the distances from proximal +, proximal -, distal + and distal - MLL2 peaks that lost H3K4me3 in double-KO mESC to the 5’UTR of the nearest L1 element. Distal - MLL2 peaks are significantly closer to L1 elements, with 62.5% of the peaks located within 1 Kb of the nearest L1 (dashed vertical line). p-values were calculated using a Wilcoxon Rank-Sum test. (D) The percentage of L1 elements belonging to the indicated L1 subfamilies is plotted for either all L1 elements present in the mouse genome (purple) or the subset of L1 elements located within 1 Kb from a distal - MLL2 peak losing H3K4me3 in double-KO mESC (yellow). The top 5 most abundant subfamilies on the distal - MLL2 peaks that lost H3K4me3 in double-KO group are shown. (E) Heatmap plots showing the GC content and the MLL2, H3K4me3, H3K27ac, H3K27me3 and H3K9me3 ChIP-seq signals in WT and double-KO mESC around the 5’UTR of L1 elements located <1 Kb from distal - MLL2 peaks losing H3K4me3 in double KO mESC (n=2763). L1s are sorted by the H3K4me3 signal in WT mESC. (F-G) Genome browser snaphots showing the MLL2, H3K4me3 and H3K9me3 ChIP-seq signals in WT and double-KO mESC at the 5’UTR of the L1 elements near (F) Smpdl3a and (G) Tcstv3 (highlighted in grey in Fig. 3A-B). The MLL2 peak found within the 5’UTR of each L1 element is highlighted in grey, showing increased H3K9me3 signal in double-KO compared to WT mESC. GC percent and Repeat Masker tracks are shown at the top. (H) Cumulative frequency distribution plot for the distances from the TSS of unaffected, upregulated and downregulated genes in double-KO mESC to the nearest MLL2-bound L1 element (i.e. L1 elements whose 5’UTR are located <1 Kb from a distal - MLL2 peak losing H3K4me3 in double-KO mESC). p-values were calculated using a Wilcoxon Rank-Sum test.

It is well established that H3K9me3, together with DNA methylation, plays a central role in repressing transposable elements (TE), including L1^62,63^. To evaluate whether MLL2 could antagonize H3K9me3-mediated repression at the identified MLL2-bound L1 elements, we performed ChIP-seq experiments for H3K9me3 in WT and double-KO ESC (Fig. 3E-G and Fig. S6C). Notably, we observed that, in double-KO ESC, the H3K9me3 mark spread towards the 5’UTRs of the MLL2-bound L1 elements, occupying the regions bound by MLL2 and enriched in H3K4me3 in WT cells. These observations further suggest that, in WT mESC, the 5’UTR/promoters of the MLL2-bound L1 elements are active but become silent in the absence of MLL2.

Several studies have demonstrated that TE, including L1, can function as enhancers during early development^81,82^. We hypothesized that if the MLL2-bound L1 elements act as enhancers whose regulatory activity is dependent on MLL2, then they could be important for the expression of genes that get downregulated in double-KO ESC. To test this hypothesis, we performed a cumulative frequency distribution analysis comparing the distances between genes that are downregulated, upregulated or unaffected in double-KO ESC with respect to the nearest MLL2-bound L1 element losing H3K4me3 in double-KO ESC (Fig. 3H). Importantly, this analysis revealed mild but significant distance differences, with the downregulated genes being closer to MLL2-bound L1 elements than the other two gene categories. Altogether these results suggest that MLL2 might influence the expression of nearby genes in ESC by controlling the regulatory activity of nearby L1 elements with enhancer-like functions.

### MLL2 controls gene induction upon ESC differentiation through L1 elements with enhancer-like activity

Most genes that are downregulated in double-KO ESC are already lowly expressed in their WT counterparts (Fig. S4B), thus questioning whether their additional silencing has any major biological impact on pluripotency. Furthermore, *Mll2^-/-^* mice do not show obvious cell type-specific defects until E9.5, suggesting that MLL2’s regulatory function is particularly relevant during differentiation, when lineage specific gene expression programs are established. Therefore, we then evaluated whether MLL2 affects gene expression during ESC differentiation and whether the MLL2-bound L1 elements contribute to proper gene induction. Briefly, the mESC lines described in previous sections (i.e. WT, *Mll1*-KO, *Mll2*-KO and double-KO) were differentiated using a four days multilineage differentiation protocol^83^ and analyzed by RNA-seq and ChIP-seq (Fig. 4A). This differentiation protocol was chosen to maximize the number of genes undergoing expression changes. In accordance with the results obtained in undifferentiated ESC (Fig. S3B), the majority of the DEG on day 4 mutant cells were downregulated, with MLL2 playing, compared to day 0, a more prominent role as reflected by the low number of DEG detected in *Mll1*-KO cells and the high and similar number of DEG identified in both *Mll2*-KO and double-KO cells (Fig. 4B; Supplementary Data 2). Importantly, almost 40% of the genes downregulated in day 4 double-KO cells corresponded to genes that got induced upon differentiation of the WT ESC (*p*-value 7.13e-63; Fig. 4C), thus indicating that MLL2 is required for proper gene induction during ESC differentiation. On the other hand, there was a significant overlap between the downregulated genes in double-KO cells on either day 0 or day 4 (Fig. S7A), with more than 1/3 of all downregulated genes on day 4 being already downregulated on day 0. These results suggest that the defects in gene induction observed upon differentiation of double-KO ESC could be attributed, at least partly, to transcriptional and/or chromatin defects already occurring in pluripotent cells. To test this possibility, we differentiated the *Mll1*-KO ESC and treated them with 4-OHT to delete *Mll2* at different timepoints (Fig. S7B): two days before the beginning of the differentiation (i.e. as in previous experiments), on the same day the differentiation was started, as well as one or two days after the beginning of the differentiation. We then assessed the expression of genes downregulated in day 4 double-KO cells by RT-qPCR (Fig. S7B). As expected, removal of *Mll2* before starting the differentiation led to the downregulation of the target genes. Interestingly, the removal of *Mll2* at later stages, including after pluripotency exit (i.e. two days after the beginning of the differentiation), also impaired the activation of the investigated genes. These results suggest that the role of MLL2 in facilitating gene induction cannot be solely attributed to its regulatory function in pluripotent cells.

**Figure 4.**
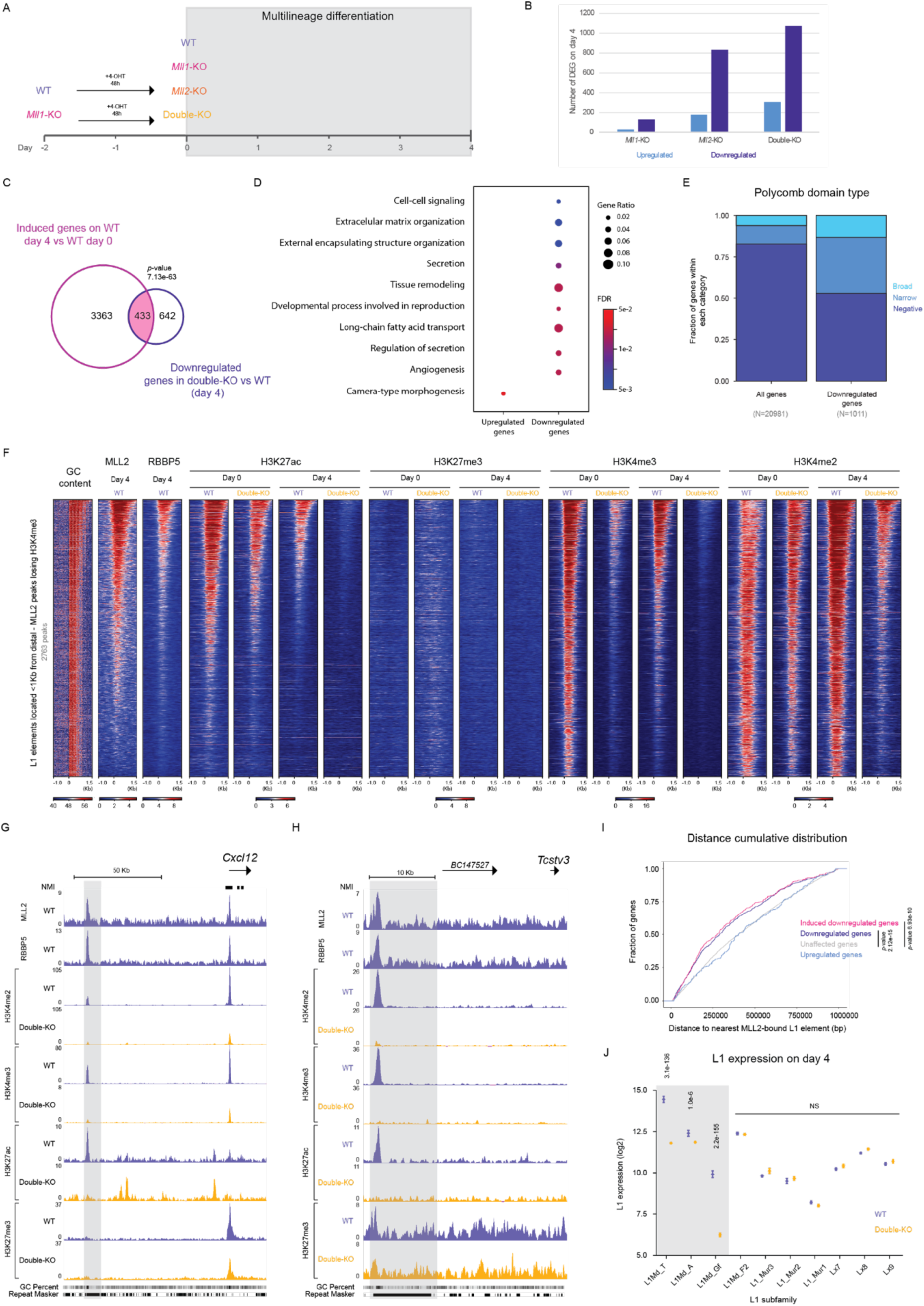
MLL2 controls gene expression upon ESC differentiation through L1 elements with enhancer-like functions. (A) Schematic representation of the experimental workffow used to investigate the regulatory function of MLL2 upon ESC differentiation. WT and Mll1-KO mESC were treated with 800 nM 4-OHT for 48 hours, after which cells were subjected to a 4-day multilineage differentiation protocol. (B) Bar plot showing the number of DEG in Mll1-KO, Mll2-KO and double-KO cells compared to WT cells on day 4. Upregulated genes are shown in blue and downregulated genes in purple. (C) Overlap between the genes that got induced upon differentiation of WT ESC (i.e. upregulated in WT day 4 vs WT day 0) and the downregulated genes in day 4 double-KO cells in comparison to day 4 WT cells (double-KO day 4 vs WT day 4). The p-value was calculated using a hypergeometric test. (D) GO enrichment analysis for the “biological process” category was performed for up-(left) and downregulated (right) genes on day 4 (double-KO vs WT) using WebGestalt^CS^. Representative examples from the top 20 most enriched terms are shown. (E) Distribution of genes according to their associated Polycomb domains (broad (≥ C Kb), narrow (0< length <C Kb) and negative (0 Kb)) is shown for either all mouse genes (left; n=20S81) or downregulated genes in day 4 double-KO cells (right; n=1011). (F) Heatmap plots showing the GC content and the MLL2, RBBP5, H3K27ac, H3K27me3, H3K4me3 and H3K4me2 ChIP-seq signals in WT and double-KO cells at the indicated differentiation time points (day 0 or day 4) around the 5’UTR of MLL2-bound L1 elements (i.e. L1 elements whose 5’UTR is located <1 Kb from a distal-MLL2 peak losing H3K4me3 in double KO mESC; n=27C3). L1s are sorted by H3K4me3 signals in WT day 4 cells. (G-H) Genome browser snapshots showing the MLL2, RBBP5, H3K4me2, H3K4me3, H3K27ac and H3K27me3 ChIP-seq signals in day 4 WT and day 4 double-KO cells around (G) Cxcl12 and (H) Tcstv3, two genes downregulated in both day 0 and day 4 double-KO cells in comparison to their corresponding WT controls. The nearby MLL2-bound L1 elements are highlighted in grey. (I) Cumulative frequency distribution plot for the distances from the TSS of unaffected, upregulated, downregulated and induced-downregulated genes in day 4 double-KO cells to the nearest MLL2-bound L1 element (i.e. L1 elements whose 5’UTR are located <1 Kb from a distal + MLL2 peak losing H3K4me3 in double-KO mESC). The “induced downregulated genes” category includes those genes that are downregulated in day 4 double KO-cells in comparison to day 4 WT cells and that, in addition, are induced upon differentiation of WT cells (i.e. upregulated in WT day 4 vs WT day 0). The p-values were calculated using a Wilcoxon Rank-Sum test (J) The expression of the indicated L1 subfamilies in WT and double-KO cells on day 4 was measured by RNA-seq using TEtranscripts^85^. For each L1 subfamily, the average expression levels and the corresponding standard deviation are plotted. The p-values were calculated using DEseq2 from TEtranscripts.

Having shown that MLL2 is important for gene induction during ESC differentiation, we then had a closer look at the type of genes whose expression was compromised in day 4 double-KO cells. The role of MLL2 in gene regulation has been previously associated with the establishment and maintenance of chromatin bivalency at the promoters of developmental genes in pluripotent cells, which in turn can prime and facilitate the subsequent activation of these genes during differentiation^11,30,84^. However, a GO analysis of the downregulated genes in day 4 double-KO cells revealed that, as already observed on day 0 (Fig. 1E), developmental terms were not particularly enriched (Fig. 4D). Consistent with the GO analysis results, the promoters of the genes downregulated in day 4 double-KO cells were mostly associated with negative or narrow H3K27me3 domains (Fig. 4E; Supplementary data 2), further supporting that, as observed in ESC (Fig. 1J), MLL2 does not preferentially control the induction of major developmental genes (i.e. promoters with broad H3K27me3 domains) during differentiation. These results also imply that antagonizing PcG-mediated repression is not the main mechanism through which MLL2 contributes to gene expression control in neither ESC nor upon their differentiation. To further test this notion and, thus, gain mechanistic insights into how MLL2 might control gene induction, we performed ChIP-seq experiments, similar to the ones described for ESC, in day 4 WT and double-KO cells. First, we analyzed those genes whose promoters were associated with broad H3K27me3 domains and that got induced during our four-days differentiation (n=627 genes; Fig. S7C), thus representing developmental genes that became induced. Overall, the promoters of these genes were bound by MLL2 in day 4 WT cells and, consistent with their transcriptional activation, gained H3K27ac and H3K4me3 and lost H3K27me3 in day 4 WT cells, compared to day 0 WT ESC. Notably, although the double-KO ESC showed lower H3K4me3 levels than WT ESC at these broad promoters, upon differentiation (i.e. day 4) the double-KO cells gained H3K4me3 and there were only subtle differences in both active (i.e. H3K4me3 and H3K27ac) and repressive (i.e. H3K27me3) marks between WT and double-KO cells. Accordingly, the majority (94%) of “broad” genes were properly induced in day 4 double-KO cells (Fig. S7D). Therefore, despite losing H3K4me3 and, thus, chromatin bivalency in double-KO ESC, the majority of developmental gene promoters get properly activated upon differentiation (i.e. gained H3K4me3 and H3K27ac). Together, these results question the functional relevance of bivalent chromatin at developmental gene promoters and suggest that other TrxG complexes might compensate for the loss of MLL2 during the activation of these promoters.

Next, given that MLL2 was bound to a large number of PE-like distal elements that might get activated, and thus, functionally relevant upon pluripotent cell differentiation^48–50^, we explored the possible involvement of these distal elements in the regulation of MLL2 target genes in day 4 cells. A cumulative frequency distribution analysis comparing the distances between genes that are downregulated, upregulated or unaffected in day 4 double-KO cells with respect to the nearest distal + MLL2 peak showed no differences between the considered gene groups (Fig. S7E). Moreover, we then identified those distal + MLL2 peaks that gained H3K27ac in day 4 WT cells (i.e. became active in day 4; n=396; Supplementary data 1) and compared their chromatin features in WT and double-KO day 4 cells (Fig.S7F). No significant differences were observed and the majority of these distal + MLL2 peaks got properly activated (i.e. gained H3K27ac) in the absence of MLL2 (Fig. S7F). Altogether, the results suggest that the regulatory function of MLL2 during gene induction does not preferentially involve the control of PE-like distal elements.

Having established a link between L1 elements and the regulation of MLL2 target genes in mESC, we next asked whether MLL2-bound L1 elements also contributed to the gene expression changes observed in day 4 double-KO cells. To do so, we focused on the set of MLL2-bound L1 elements identified in mESC (Fig. 3E; Supplementary Data 1) and compared their chromatin features in both WT and double-KO cells at day 0 and day 4 (Fig. 4F-H). In day 4 WT cells, MLL2 and RBBP5 remained bound to the 5’UTR of a significant fraction of these L1 elements, and this binding was accompanied by the retention of H3K27ac and H3K4me3, suggesting that at least some of the MLL2-bound L1 elements remained active upon differentiation (Fig. 4F-H). Interestingly, for those L1 elements still bound by MLL2 on day 4, H3K27ac and H3K4me3 were almost completely lost in day 4 double-KO cells (Fig. 4F-H), suggesting that their activity becomes increasingly dependent on MLL2 during differentiation. Supporting a functional role for these L1 elements, downregulated genes in day 4 double-KO cells - including those normally induced during our differentiation - were significantly closer to MLL2-bound L1 elements than either upregulated or unaffected genes (Fig. 4I). Furthermore, the distance differences between the downregulated and unaffected genes in day 4 cells were considerably more pronounced than in ESC (Fig. 4I vs Fig. 3H), suggesting that the regulatory effects of the MLL2-bound L1 elements become more pronounced upon ESC differentiation.

To further investigate the functional role of MLL2 at L1 elements, we analyzed transcriptional changes (double-KO vs WT) across different L1 subfamilies at both day 0 and day 4 (Fig. S7G, Fig. 4J). Despite the chromatin changes previously observed at the 5’UTR of the MLL2-bound L1 subfamilies on day 0 (Fig. 3E), no corresponding differences in their transcriptional level were detected (Fig. S7G). On day 4, however, the expression of three young L1 subfamilies — L1Md_T, L1Md_A and L1Md_Gf — was significantly reduced in double-KO cells compared to WT (adjusted *p*-values of 3.1e-136, 1.0e-6 and 2.2e-155, respectively). Notably, these are the same subfamilies with GC-rich 5’UTRs^51^ we previously found to be enriched in distal MLL2 peaks and that lost H3K4me3 in double-KO cells (Fig. 3D-E). Furthermore, the pronounced transcriptional changes observed for the MLL2-bound L1 elements in day 4 double-KO cells is also in agreement with the stronger association between these L1 elements and the genes downregulated in day 4 double-KO cells compared to genes downregulated in day 0 double-KO cells (Fig. 3H, Fig. 4I). Altogether, these results support the notion that MLL2 regulates the enhancer-like activity of L1 elements, which in turn can influence the expression of nearby genes in ESC and, more prominently, during their differentiation.

### MLL2 supports the long-range communication between L1 elements and their associated genes

MLL2 has been studied for its potential contribution to 3D genome architecture using a range of chromatin conformation techniques^30,84^. These studies suggest that while MLL2 has minimal impact on global chromatin organization, it does contribute locally to 3D chromatin topology, particularly at bivalent gene promoters^30,84^. MLL2 loss was described to lead to modest A-to-B compartment switching, especially in regions enriched for bivalent genes, causing a redistribution of promoter-centered interactions^30^. However, absence of MLL2 did not alter topologically associating domain (TAD) boundaries or size, neither in pluripotent cells nor during their differentiation^30,84^.

To investigate whether, despite not having a major role in global 3D chromatin organization, MLL2 could facilitate the physical communication between MLL2-bound L1 elements and nearby MLL2 target genes, we performed micro-C analysis on WT and double-KO cells at day 0 and day 4. We first assessed the impact of MLL2 loss on A/B compartmentalization (Fig. 5A-B, Fig. S8A). In agreement with previous findings^30,84^, MLL2 loss had an overall minor impact in A/B compartment organization. Nevertheless, A-to-B compartment switches were slightly more frequent than B-to-A switches in double-KO cells in comparison to WT cells (Fig. 5A), suggesting a subtle bias towards chromatin compaction upon MLL2 loss. Similarly, analysis of TAD organization revealed no significant alterations in the absence of MLL2 (Fig. 5C-D, Fig. S8B). These findings confirm that MLL2 does not play a major role in higher-order chromatin organization.

**Figure 5.**
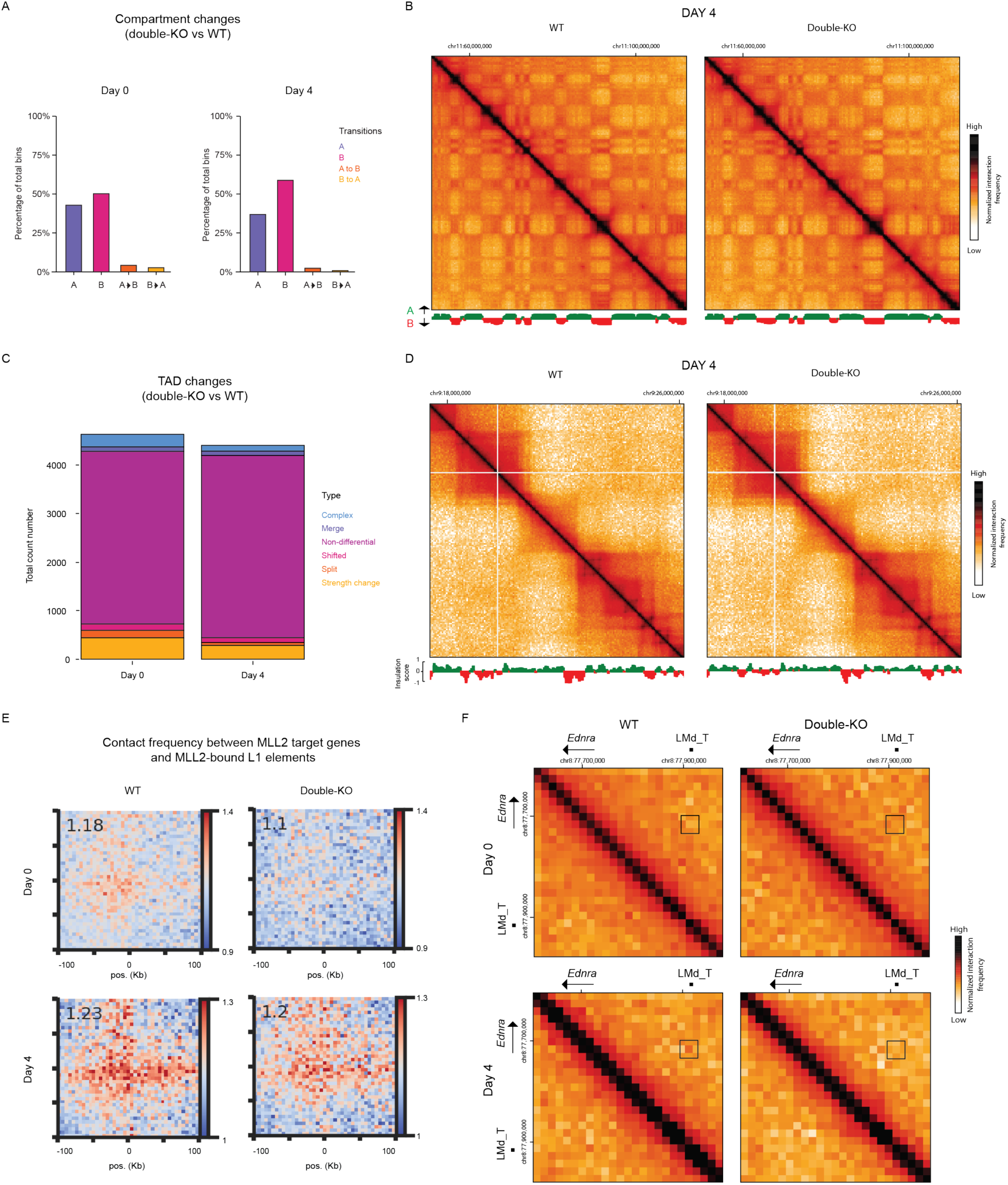
The MLL2-bound L1 elements show weak interactions with nearby target genes. (A) Bar plots displaying the percentage of compartment transitions (WT vs double-KO) on day 0 (left) and day 4 (right). Four different conditions were considered: unchanged A compartments (purple), unchanged B compartments (pink), A-to-B transitions (orange) and B-to-A transitions (yellow). (B) Micro-C normalized contact matrix for WT (left) and double-KO (right) day 4 cells at 350 Kb resolution. Eigenvector values are shown at the bottom (positive/green values correspond to A compartments and negative/red values to B compartments). (C) Distribution of TAD changes (WT vs double-KO) on day 0 and day 4 (see Methods for details). (D) Micro-C normalized contact matrix for WT (left) and double-KO (right) day 4 cells at 50 Kb resolution. Insulation score values are shown at the bottom. (E) Pileup plots of micro-C contact frequencies between pairs of MLL2-bound L1 elements and the promoters of nearby MLL2 target genes (i.e. genes whose promoter is bound by MLL2 in WT mESC and that lost H3K4me3 in double-KO mESC; see Methods for details). (F) Micro-C normalized contact matrices (15 Kb resolution) are shown around Ednra, a gene downregulated in day 4 double-KO cells. Matrices for WT (left) and double-KO (right) cells both at day 0 (top) and day 4 (bottom) are shown. The black squares highlight the contact between the Ednra promoter and a nearby MLL2-bound L1 element for each genotype and differentiation timepoint.

Our previous data suggests that MLL2 controls gene expression, especially upon ESC differentiation, by regulating the activity of a subset of L1 elements with enhancer-like regulatory properties. To further test this possibility, we performed pileup analyses of the micro-C data to evaluate whether the MLL2-bound L1 elements physically interacted with their putative target genes (Supplementary Data 1)^86^. Importantly, we found that in WT cells at day 0, these L1 elements already engaged into weak, albeit detectable, physical interactions with the promoters of nearby target genes, and these interactions were intensified on day 4 (Fig. 5E-F). Notably, in double-KO cells, these L1-target gene contacts were slightly decreased at both timepoints (Fig. 5E-F). It is worth mentioning that the overall weakness of the L1-promoter contacts in both WT and KO cells could be partly explained by the difficulties in mapping micro-C reads within repetitive elements. Taken together, these findings further support the long-range regulatory activity of the MLL2-bound L1 elements and suggest that MLL2 contributes to, but it is not essential for, the physical communication between L1 elements and nearby target genes.

### Deletion of representative MLL2-bound L1 elements demonstrates that they can act as enhancers of nearby MLL2 target genes

The previous results provide strong correlative observations implicating MLL2-bound L1 elements in the long-range regulation of MLL2 target genes both in ESC as well as upon their differentiation. To directly assess whether the MLL2-bond L1 elements act as enhancers of MLL2 target genes, we used CRISPR-Cas9 to delete the L1 elements near three genes that were downregulated in double-KO cells in both day 0 and day 4 and whose promoters were associated with different types of Polycomb domains (Fig. 6 and Fig. S9A-D): *Cxcl12* (broad), *Cnrip1* (narrow) and *Tcstv3* (negative). Once several clonal mESC lines with homozygous deletions of the selected L1 elements were obtained (five clones for *Cxcl12*, four clones for *Cnrip1* and four clones for *Tcstv3*), we measured the expression of the associated genes by RT-qPCR in both day 0 and day 4 (Fig. 6A-C). Importantly, in all three cases, the deletion of the MLL2-bound L1 element led to the downregulation of the nearby MLL2 target gene at both day 0 and day 4 (Fig. 6A-C). These results demonstrate that the L1 elements whose promoters are bound and regulated by MLL2 can indeed act as enhancers of the genes downregulated in double-KO cells.

**Figure 6.**
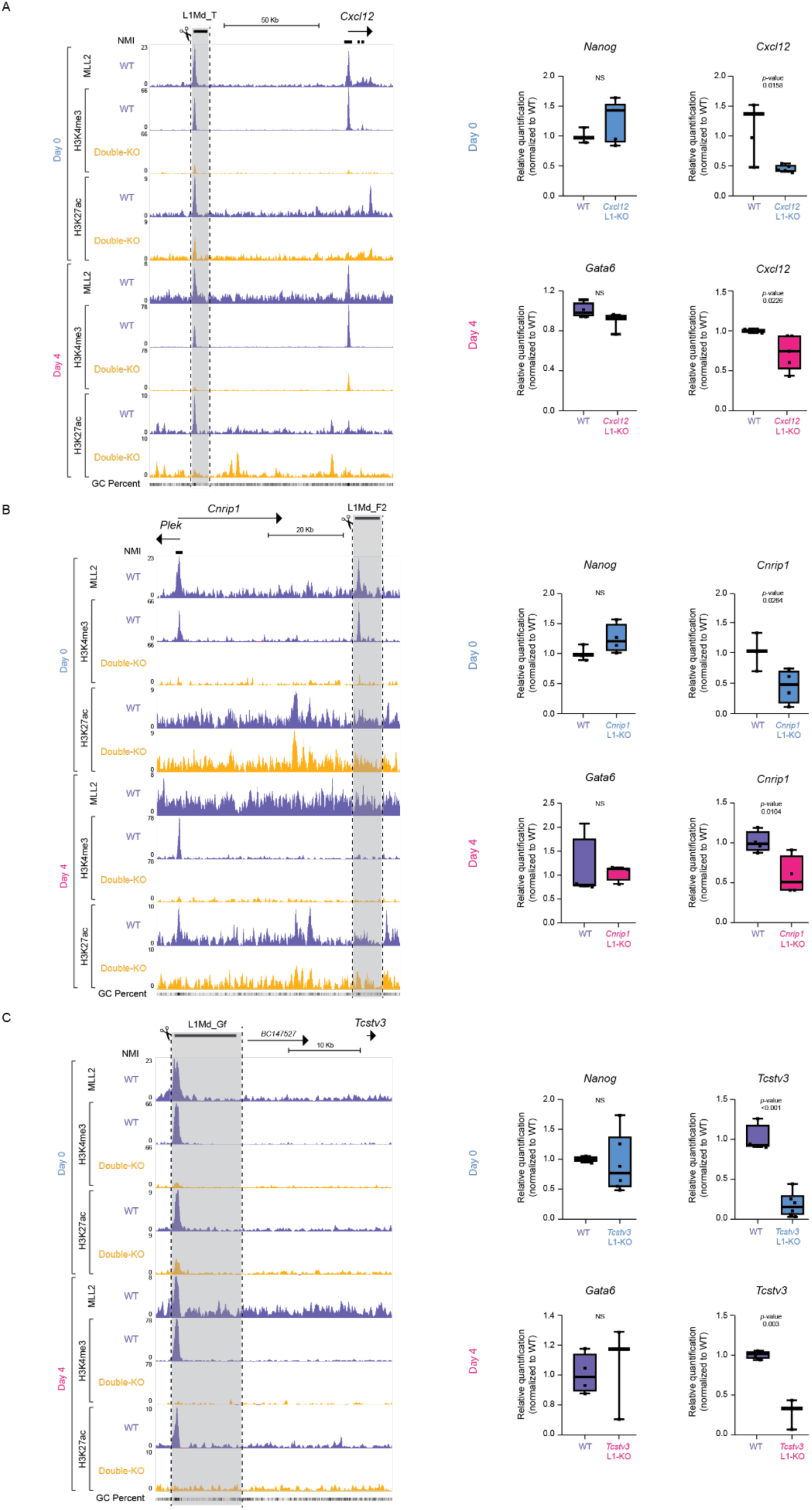
Deletion of MLL2-bound L1 elements leads to downregulation of nearby target genes. (A-C) Genome browser snapshot highlighting the MLL2-bound L1 elements located at the (A) Cxcl12, (B) Cnrip1 and (C) Tcstv3 loci and that were deleted by CRISPR-Cas9 genome editing. For each L1 element deletion, the expression levels of the corresponding target gene, as well as the expression of Nanog or Gata6 (used as controls for day 0 and day 4, respectively) were measured by RT-qPCR on day 0 and 4 in WT cells and cells with homozygous deletions of each MLL2-bound L1 element. For each time point (i.e. day 0 or day 4), the expression values were normalized to WT cells using Hprt and Eef1a1 as loading controls. The following number of replicates were analyzed for each gene: (A) for Cxcl12, three biological replicates of WT cells and five Cxcl12 L1-KO clones (one biological replicate per clone) were analyzed on day 0; four biological replicates of WT cells and three and five Cxcl12 L1-KO clones (one biological replicate per clone) for Gata6 and Cxcl12 RT-qPCR, respectively, were analyzed on day 4; (B) for Cnrip1, three biological replicates of WT cells and four Cnrip1 L1-KO clones (one biological replicate per clone) were analyzed on day 0; four biological replicates of WT cells and four Cnrip1 L1-KO clones (one biological replicate per clone) were analyzed on day 4; (C) for Tcstv3, four biological replicates of WT cells and four Tcstv3 L1-KO clones (one biological replicate per clone) were analyzed on day 0; four biological replicates of WT cells and three Tcstv3 L1-KO clones (one biological replicate per clone) were analyzed on day 4. The p-values were calculated using unpaired t-tests. NS: not significant.

## Discussion

Understanding how genetic and epigenetic mechanisms interact to regulate gene expression remains a fundamental question in chromatin biology. Bivalent promoters — characterized by the coexistence of active and repressive chromatin marks — represent a key example of this complexity^11,87–89^. At these loci, TrxG and PcG complexes come together, establishing unique chromatin signatures that are thought to poise developmental genes for future activation^29,30,90^. MLL2 is the primary methyltransferase responsible for establishing and maintaining H3K4me3 at bivalent promoters in ESC^29–31,90,91^. Moreover, the loss of MLL2 in ESC results in downregulation of a subset of genes partially overlapping with those marked by bivalent promoters^30^. Consequently, it has been proposed that, in pluripotent cells, MLL2 plays a critical regulatory role at bivalent promoters by counterbalancing the repressive activity of PcG proteins and keeping them poised for activation, therefore, preserving the epigenetic plasticity necessary for differentiation and proper embryonic development^30^. Previous work suggesting that promoter bivalency does not confer faster gene induction dynamics, together with the identification of bivalent chromatin in differentiated cells, is now challenging this view and questioning the functional relevance of promoter bivalency during development^22,23,92^. Additionally, inconsistencies between the genes that lose H3K4me3 and those that become downregulated upon loss of MLL2, together with discrepancies regarding the changes in PcG binding levels at bivalent promoters in *Mll2*^-/-^ cells^30,44^, led to an ongoing discussion over the mechanism through which MLL2 regulates gene expression in pluripotent cells. In agreement with previous reports^30^, we confirmed that MLL2 is required in ESC for the maintenance of H3K4me3 at the bivalent promoters of major developmental genes. However, our data shows that for the majority of these promoters, the loss of H3K4me3 in ESC does not affect gene expression (Fig. 1D,J). Accordingly, we observed only mild increases in H3K27me3/PRC2 levels and no changes for PRC1 at MLL2-dependent bivalent promoters (Fig. 2B). Given that PRC1, rather than PRC2, is the main responsible for PcG-mediated repression in ESC^77,78,93^, our results suggest that counteracting PcG-mediated repression at developmental gene promoters is not the main regulatory function of MLL2 in pluripotent cells. Furthermore, bivalency loss in *Mll2*-null ESC did not substantially impair the activation of key developmental genes upon differentiation (Fig. S7C-D), probably due to the compensatory effects of other TrxG complexes at active promoters^94^. In addition, approximately half of the genes regulated by MLL2 in either ESC or upon differentiation are not PcG targets, as their promoters are not enriched in H3K27me3 (Fig 1J and Fig 4E), suggesting that MLL2 might control their expression through mechanisms other than antagonizing PcG-mediated repression. Overall, our findings question the functional relevance of promoter bivalency during pluripotent cell differentiation and indicate that, during this process, MLL2 might preferentially control gene expression through PcG-independent regulatory mechanisms. Nevertheless, the developmental relevance of the antagonism between TrxG and PcG might be context dependent and particularly important in specific cell lineages^95^ or developmental loci (e.g. *Hox* clusters)^42^.

Having shown that antagonizing PcG-mediated repression is not the main regulatory function of MLL2 in neither pluripotent nor differentiating cells, we then explored alternative mechanisms through which MLL2 could regulate gene expression. Chiefly, here we show that MLL2 actively binds to and regulates young L1 elements with enhancer-like activity, which in turn significantly contribute to the regulation of MLL2 target genes in ESC and, especially, upon their differentiation. TE, such as L1, are known to evade repression in the host cells at various stages of embryonic development, including in cultured ESC (as reviewed by ^96^). Expression of TE during development is not a simple consequence of increased chromatin openness as several individual TE families have been found to be essential for the correct progression of embryonic development^97,98^, which, among other mechanisms, often involves long-range regulation of nearby genes (i.e. enhancer-like activity)^57,66,81,99^. In recent years, work from several laboratories has highlighted the importance of L1-mediated gene regulation during development, identifying various regulators of L1 activity as necessary for proper gene expression control^61,66,81,100^. These regulators include specific histone PTMs^81^, RNA polII elongation factor 3 (ELL3)^66^, enzymes involved in RNA metabolism^100^ and the Human Silencing Hub (HUSH) complex^61^. Together, these findings suggest the existence of a complex multilayered network involved in gene regulation through L1 elements that is not yet fully understood. A previous study reported that the repressor ZFP281 and MLL2 bind to the 5’UTR of young L1 subfamilies (e.g. L1Md_A, L1Md_T) in mESC, albeit with opposing binding patterns^52^. Regarding their function, ZFP281 represses the activity of these young L1 elements by recruiting Tet1 and Sin3A to their 5’ UTR^52^, thereby restricting MLL2 access. Our data provides compelling evidence that not only supports the essential role of MLL2 in controlling the activity of young L1 subfamilies in mESC, but also upon their differentiation. Most importantly, we also show that these MLL2-dependent L1 elements display enhancer-like activity and are required for the proper expression of neighboring MLL2-target genes, both in pluripotent and differentiating cells.

MLL2 binds to and regulates the 5’UTR/promoter of a discrete set of young L1 elements belonging to specific subfamilies (e.g. L1Md_A, L1Md_Gf and L1Md_T). Interestingly, the 5’UTR of these MLL2-bound L1 elements share a rather unique genetic composition consisting of CG-rich sequences resembling CGI, which are probably not classified as such due to their repetitive nature (Fig. 3E). The presence of these CG-rich sequences could explain how MLL2 COMPASS-like complexes are specifically targeted to the L1 5’UTRs through the MLL2 CXXC domain^71^. However, there are other proteins besides MLL2 that also contain CXXC domains recognizing non-methylated CGI (e.g. MLL1, CFC1, KDM2B)^71,101^ that nevertheless, do not seem to be recruited to the 5’UTR of the MLL2-bound L1 elements: (i) the MLL2-bound L1 5’UTRs are not enriched in H3K27me3 regardless of whether they are active (i.e. marked with H3K27ac) or not (Fig. 3E), suggesting that KDM2B, a component of PRC1 responsible of the recruitment of PcG complexes to CGI, cannot recognize the L1 5’UTR sequences; (ii) SET1A/B complexes are largely responsible of H3K4me3 at CGI-rich active promoters in ESC^94,102,103^, explaining why the loss of MLL2 does not affect H3K4me3 levels at such locations (Fig 1B). In contrast, H3K4me3 levels at the MLL2-bound L1 5’UTRs are severely reduced in the absence of MLL2, regardless of whether the 5’UTR are active (i.e. marked with H3K27ac) or not. This suggests that CFP1, the CXXC protein targeting SET1A/B complexes to active CGI^102^ cannot properly recognize the MLL2-bound L1 5’UTR sequences; (iii) in the absence of MLL2, the binding of MLL1 strongly increases at “canonical” CGI that are either active or bivalent (Fig 1B-C). However, this compensatory mechanism is not observed at the MLL2-bound L1 5’UTRs, which show low MLL1 binding levels in both WT and *Mll2*-KO ESC (Fig. S6B). It is currently unclear why MLL2, and not other CXXC proteins, can bind to the CG-rich L1 5’UTR sequences, although it is tempting to speculate that it could involve differences in DNA-binding selectivity among CXXC domains as well as the presence of additional DNA-binding domains unique to each CXXC protein^101,104^. Regardless, our data suggest that the binding to the CG-rich 5’UTR sequences of young L1 elements might be a unique feature of MLL2 among CXXC proteins, thus making these 5’UTR/promoters particularly dependent on MLL2 COMPASS-like complexes. Finally, our work does not provide a direct mechanistic explanation as to how MLL2 controls the activity of the L1 5’UTR/promoters and, thus, of nearby MLL2 target genes. Recent work addressing the function of H3K4me3 at promoter regions indicates that the presence of this histone mark, which at L1 5’UTR/promoters is fully dependent on MLL2, might increase productive elongation by facilitating RNA polII pause release and/or preventing premature termination^10,104,105^. In addition, our data suggest that the presence of MLL2 at the 5’UTR of young L1 elements prevents the spreading of H3K9me3, which is particularly important for the silencing of young L1 elements in both mESC and hESC^62,63,106^. One potential mechanism through which MLL2 could protect the L1 5’UTR/promoters from H3K9me3 and, thus, keep them active, is by recruiting the KDM4A/C H3K9me3 demethylases through H3K4me3^63,107^. Last but not least, previous studies implicated MLL2 in the control of 3D genome organization, albeit the observed changes in chromatin architecture were rather minor^30,82,84^. In agreement with this, we found that although the MLL2-bound L1 elements interact with nearby MLL2 target genes, thus further supporting their enhancer-like function, such interactions seem to be weak and only mildly affected in the absence of MLL2.

Overall, our findings shed light on a poorly explored role of MLL2 in regulating gene expression through young L1 elements with enhancer-like activity, thus expanding MLL2 function beyond modulating PcG-mediated repression at bivalent promoters in both pluripotent and differentiating cells. Understanding how MLL2 is targeted to these L1 elements, how these L1 elements are capable of long-range expression control and how this regulation might change depending on the cellular context remain as key questions that, if solved, will provide important insights into the broader roles of bivalency, transposable elements, and epigenetic plasticity during embryogenesis.

## Supporting information

Supplementary data description

## Author contributions

LZS and ARI conceptualized and designed the study. LZS carried out the experimental work. LDF and SMR performed data analyses. LZS, LDF and ARI wrote the manuscript. ARI supervised the experiments and data analyses.

## Acknowledgements

We thank F. Stewart (TU Dresden) for kindly providing the Cre-inducible *Mll2*-KO mESC line used in this study. We would like to thank all the Rada-Iglesias lab members for insightful comments and suggestions. Work in the Rada-Iglesias laboratory is supported by the following grants: PID2021-123030NB-I00 (ARI), funded by MCIN/AEI/10.13039/501100011033 and by ERDF “A way of making Europe”; RED2022-134100-T (ARI) (REDEVNEURAL 3.0), funded by MCIN/AEI/10.13039/501100011033; ERC Consolidator Grant Poised-Logic (862022) (ARI), funded by the European Research Council; ENHPATHY H2020-MSCA-ITN-2019-860002 (ARI), funded by the European Commission; and Chromrare HORIZON-MSCA-2021-DN-01-101073334 (ARI), also funded by the European Commission. LZS was supported by an EMBO Postdoctoral fellowship.

## Material G Methods

### Experimental Methods

#### Cell culture

mESC were grown on 0.1% gelatin-coated plates, using KnockOut^TM^ DMEM (Thermo Fisher Scientific, 10829018) supplemented with 15% fetal bovine serum (FBS; Thermo Fisher Scientific, A5256801), 1x antibiotic and antimycotic solution (Sigma-Aldrich, A5955), 1x glutaMAX (Thermo Fisher Scientific, 35050038), 1x non-essential amino acids (NEAA; Thermo Fisher Scientific, 11140035), 0.1 mM β-mercaptoethanol (Thermo Fisher Scientific, 21985023) and leukemia inhibitory factor (LIF; made in-house). Cells were maintained at 37 °C with 5% CO_2_. For the multilineage differentiation, cells were grown on 0.1% gelatin-coated plates, using KnockOut^TM^ DMEM supplemented with 10% FBS, 2 mM L-glutamine (Thermo Fisher Scientific, 25030024), 0.1 nM β-mercaptoethanol, 1x NEAA, 1x antibiotic and antimycotic solution and 1 µM retinoic acid (RA; Sigma-Aldrich, R2625) for 4 days, as previously described by ^83^.

#### CRISPR-CasG genome editing and genotyping of transgenic mESC lines

For the generation of *MLL1*-KO, *Cxcl12* L1-KO, *Cnrip1* L1-KO and *Tcstv3* L1-KO cell lines, two sg-RNAs were designed flanking the region to-be deleted for each line, using Benchling’s CRISPR tool (https://www.benchling.com/crispr). For each sgRNA, two oligonucleotides were synthesized (Supplementary Data 3), annealed and cloned into a CRISPR-Cas9 expression vector (pX330-hCas9-long-chimeric-grna-g2p, provided by Leo Kurian’s lab). mESC were transfected with CRISPR-Cas9 constructs using Lipofectamine^TM^ 3000 (Thermo Fisher Scientific, L3000001). After 16-24 hours, cells were subjected to puromycin selection for 48 hours. Surviving cells were single cell-plated in 96-well plates through serial dilutions. Individual clones were screened for the intended deletion via PCR (NZYTaq II 2× Green Master, NZYtech, MB35803) with the primers listed in Supplementary Data 3, and positive clones were further validated by Sanger sequencing.

#### 4-OHT and GSK126 treatment

For the generation of *Mll2*-KO and double-KO mESC lines, cells were treated with (Z)-4-Hydroxytamoxifen (4-OHT; Sigma-Aldrich, H7904) at a final concentration of 800 nM for 48 hours. The efficiency of the 4-OHT treatment was assessed after each experiment by PCR using GoTaq® DNA Polymerase (Promega, M3005). The sequence of the primers used can be found in Supplementary Data 3.

For PRC2 inhibition assays, mESC were incubated with GSK126 (MedChemExpress, HY-13470), at a final concentration of 5 µM for 72 hours. Treatment with GSK126 on *Mll2*-KO and double-KO cells was carried out immediately following 4-OHT treatment.

#### Protein extraction and western blotting

To isolate histones, cell pellets were first incubated for 10 minutes in TEB buffer (containing 0.5% Triton X-100 in PBS) to obtain nuclei. Histones were then extracted by overnight (O/N) incubation at 4 °C in 0.2 N HCl.

To extract total proteins, plated cells were first washed twice with cold PBS (137 mM NaCl, 2.7 mM KCl, 10 mM Na₂HPO₄, 1.8 mM KH₂PO₄). RIPA buffer (0.2% SDS, 1% Triton X-100, 1 mM EDTA, 150 mM NaCl, 50 mM Tris-HCl pH 8.0, 0.5% sodium deoxycholate) was then added directly to the cells. The cells were scraped off and transferred into 1.5 mL microcentrifuge tubes. Samples were incubated on a rotating wheel at 4 °C for 30 minutes, followed by brief sonication (2 minutes, at 50% amplitude, 30 seconds ON 30 seconds OFF). Afterwards, samples were centrifuged at 14000 rpm for 15 minutes at 4 °C. The resulting supernatant was collected, aliquoted into fresh 1.5 mL tubes, and either used immediately or stored at −80 °C for future use.

Following extraction, protein concentrations were determined using the Bradford assay. Equal protein quantities were loaded onto SDS-PAGE gels, separated by electrophoresis, and transferred onto nitrocellulose membranes. The antibodies used are listed on Supplementary Data 4.

#### ChIP-seq

Approximately 10 million and 40 million cells were used for histone PTMs and COMPASS subunits ChIPs, respectively. For histone modification ChIP assays, cells were crosslinked with 1% formaldehyde for 10 minutes at room temperature (RT) before being quenched with 0.125 M glycine for another 10 minutes, while for COMPASS subunits ChIP assays cells were double crosslinked with ChIP cross-link Gold (Diagenode, C01019027) and 1% formaldehyde according to the manufactor’s instructions. Chromatin was isolated through sequential incubation with three distinct lysis buffers: LB1 (50 mM HEPES-KOH pH 7.5, 140 mM NaCl, 1 mM EDTA, 10% glycerol, 0.5% NP-40, 0.25% Triton X-100), LB2 (10 mM Tris-HCl pH 8.0, 200 mM NaCl, 1 mM EDTA, 0.5 mM EGTA), and LB3 (10 mM Tris-HCl pH 8.0, 100 mM NaCl, 1 mM EDTA, 0.5 mM EGTA, 0.1% sodium deoxycholate, 0.5% N-lauroylsarcosine).

Chromatin sonication was carried out in LB3 using an EpiShear^TM^ sonicator (Active Motif; 4-5 minutes, 20 seconds ON and 30 seconds OFF at 25% amplitude). The sheared chromatin was then incubated O/N at 4 °C with either 5 or 10 μg of antibody for histone PTMs or COMPASS subunits respectively (the list of antibodies used can be found in Supplementary Data 4). For each ChIP, 100 μL of prewashed (0.5% BSA in 1× PBS) Dynabeads (50% protein G (Thermo Fisher Scientific, 10004D) 50% protein A Dynabeads (Thermo Fisher Scientific, 10002D)) were used to capture antibody-bound chromatin during a 4-hour incubation at 4 °C. Subsequently, the beads were washed five times with RIPA buffer (50 mM HEPES-KOH pH 7.5, 500 mM LiCl, 1 mM EDTA, 1% NP-40, and 0.7% sodium deoxycholate) before being eluted (50 mM Tris-HCl pH 8.0, 10 mM EDTA and 1% SDS). The samples were then reverse crosslinked at 65 °C, O/N. The samples were treated with 0.2 mg/mL RNase A (Thermo Fisher Scientific, EN0531) and 0.2 mg/mL proteinase K (Thermo Fisher Scientific, EO0491), and finally, the DNA was purified using the QIAquick PCR Purification Kit (Qiagen, 28104).

#### RNA, cDNA synthesis and RT-qPCR

Total RNA was extracted using the NZY Total RNA Isolation Kit (NZYTech, MB13402), according to the protocol provided by the manufacturer. Complementary DNA (cDNA) was synthesized from the isolated RNA using the ProtoScript® II First Strand cDNA Synthesis Kit (New England Biolabs, E6560L). For each reaction, 1 µg of RNA was used.

RT-qPCR was conducted using the CFX 384 Real-Time PCR Detection System (Bio-Rad), employing the 2x NZYSpeedy qPCR Green Master Mix (NZYtech, MB224). Each sample was analyzed in technical triplicates using the primers detailed in Supplementary Data 3. The number of biological replicates and independent clonal lines per cell line is specified in the respective figure legends.

Relative expression levels were determined using the 2^−ΔΔCT^ method, normalizing to *Eef1a1* and *Hprt1*, two housekeeping genes that were used as loading controls.

#### Micro-C

The micro-C library was prepared using the Dovetail® Micro-C Kit according to the manufacturer’s protocol. Briefly, the chromatin was fixed with disuccinimidyl glutarate (DSG) and formaldehyde in the nucleus. The cross-linked chromatin was then digested *in situ* with micrococcal nuclease (MNase). Following digestion, the cells were lysed with SDS to extract the chromatin fragments, and the chromatin fragments were bound to Chromatin Capture Beads. Next, the chromatin ends were repaired and ligated to a biotinylated bridge adapter followed by proximity ligation of adapter-containing ends. After proximity ligation, the crosslinks were reversed, the associated proteins were degraded, and the DNA was purified then converted into a sequencing library using Illumina-compatible adaptors. Biotin-containing fragments were isolated using streptavidin beads prior to PCR amplification. For each cell type (e.g. WT Day 0), micro-C samples were prepared as two biological replicates, and for each replicate three libraries were prepared. Each library was sequenced on an Illumina Novaseq6000 platform to generate 300 million 2×150 bp read pairs/library.

### Computational Methods

#### ChIP-seq analysis

ChIP-seq data were processed using a customized pipeline based on nf-core/chipseq v2.0.0 (10.5281/zenodo.1400710), part of the nf-core community-curated workflow collection^108^. The pipeline was executed using Nextflow v23.04.4. FASTQ files were trimmed with Trim Galore v0.6.7 (using cutadapt v3.4). Alignments to the Mus musculus mm10 (GRCm38) reference genome were performed with STAR^109^. To improve read mappability in repetitive regions, STAR v2.6.1d was used with modified mapping parameters (“mouse random mode”), as described by Teissandier et al.^68^.

Please note that for the histone mark ChIP-seq experiments, the chromatin from the mouse cells was initially mixed with chromatin from human HEK293 cells in a 90%:10% proportion for normalization purposes^110^. However, this normalization strategy was discarded due to high variability in the number of human reads obtained for each sample. Consequently, reads were aligned to a custom hybrid genome composed of mm10 and hg19 assemblies and only those reads mapping exclusively to mm10 were considered for downstream analyses^111^.

Once read mapping was concluded, the resulting BAM files were processed with samtools v1.15.1^112^ and Picard v2.27.4 (https://github.com/broadinstitute/picard) for sorting, indexing, and duplicate marking. For visualization purposes, bigwig files from biological ChIP-seq replicates were merged using deepTools v3.5.5^113^ to generate average bigwig signal tracks. DeepTools v3.5.5 was also used to generate ChIP-seq heatmap plots.

Peak calling was performed using MACS2 v2.2.7.1^114^ with the --broad option, an FDR cutoff of 0.05, and a fold enrichment threshold of 3. Differential H3K4me3 peaks between double knockout and wild-type samples were identified using DiffBind v3.14^115^, with an FDR < 0.05 and |fold change| ≥ 2 (Supplementary Data 1).

MLL2 peaks were classified in four groups (proximal +, proximal -, distal + and distal -) based on their distance to TSS and NMI^116^ according to the following criteria (Supplementary Data 1):

- Proximity to TSS: peaks within 5 Kb of a TSS were considered “proximal”; peaks located ≥5 kb from a TSS were considered “distal”.

- Overlap with NMIs: peaks within 1 Kb of an NMI were labelled as “+” and all others were labelled as “-”.

#### RNA-seq analysis

RNA-seq data were processed using nf-core/rnaseq v3.14.0 (with the --aligner star_rsem option)^108^ (10.5281/zenodo.1400710), using the mm10 (GRCm38) reference genome. The pipeline was executed with Nextflow v23.04.4. Reads were trimmed with Trim Galore 3.4, aligned with STAR 2.7.10a^109^, and quantified using RSEM 1.3.1^117^.

Differential expression analysis was performed using nf-core/differentialabundance v1.5.0, implementing DESeq2 v1.34.0 (sfType: ratio; fitType: parametric; test: Wald; altHypothesis: greaterAbs; pAdjustMethod: BH; alpha: 0.1). Differentially expressed genes were defined with FDR ≤ 0.05 and |log2FC| ≥ 1^118^ (Supplementary Data 2).

Gene ontology enrichment for the “biological process” category was performed using WebGestalt^69^, and transcription factor enrichment was assessed with Enrichr using the “ChEA 2022” option^70^.

#### Micro-C analysis

Micro-C data were processed using a custom pipeline implemented in Nextflow v23.04.4, based on the Dovetail Genomics protocol (https://micro-c.readthedocs.io/en/latest/). Reads were aligned with BWA-MEM^119^. Valid ligation events were identified using pairtools parse (--min-mapq 40, -- walks-policy 5unique, --max-inter-align-gap 30) and duplicates removed using pairtools dedup. BAM files were sorted and indexed using samtools.

Contact matrices were generated using cooler cload pairix (PMID: 31290943). For compartment analysis, 350 kb resolution matrices were used and analyzed with FAN-C^120^ on ICE-normalized data. Principal components correlating with epigenomic marks were selected. Compartment dynamics were quantified using in-house scripts.

For TAD detection, 50 Kb resolution matrices were used. Insulation scores were computed with FAN-C (multiple window sizes), and TAD boundary changes were detected using TADCompare^121^.

#### Additional computational analyses

##### Classification of genes based on H3K27me3 domains

Genes were classified into three categories based on H3K27me3 ChIP-seq data from wild-type mESC^51,73^. Adjacent H3K27me3 peaks within 1 kb were merged into domains. Domains overlapping a gene TSS were used for classifying genes based on the width of the associated H3K27me3 domains:

- Broad: >6000 bp
- Narrow: ≤6000 bp
- Negative: no detectable H3K27me3 at the TSS.

##### Cumulative frequency analyses

Cumulative distributions of distances between genomic features were computed using custom scripts. For each pair, the distance from a primary feature to the nearest secondary feature was calculated. The following analyses were performed:

- Each of the differentially expressed gene categories in double-KO cells (downregulated, upregulated, unaffected) to the nearest distal + MLL2 peak losing H3K4me3 in double-KO cells.
- Each of the four categories of MLL2 peaks losing H3K4me3 in double-KO cells (proximal +, proximal -, distal +, distal -) to the 5’UTR of the nearest L1 element.
- Each of the differentially expressed gene categories in double-KO cells (downregulated, upregulated, unaffected) to the nearest MLL2-bound L1 element.

Statistical comparisons between groups were performed using the Wilcoxon rank-sum test.

##### Contacts between MLL2-bound L1 elements and MLL2 target genes

To test whether MLL2-bound L1 elements (i.e. L1 elements whose 5’UTR was <1 Kb from a distal – MLL2 peak losing H3K4me3 in double-KO cells) were in physical proximity with MLL2 target genes, we integrated ChIP-seq and Micro-C data.

MLL2-bound L1 elements were linked to genes using GREAT^122^. Proximal MLL2 peaks losing H3K4me3 in double-KO cells were also linked to genes using GREAT and the resulting gene list was used as the “MLL2 target genes” in these analyses. Shared target genes between the two sets were used to obtain a list of “MLL2-bound L1 – MLL2 target gene” pairs (Supplementary Data 1). Those pairs were then used for pile-up analyses of the Micro-C data using coolpup.py^86^ on 5 Kb resolution matrices.

##### L1 subfamily distribution

L1 subfamilies were analyzed using RepeatMasker annotations for the mm10 genome (UCSC Genome Browser)^123^. The MLL2-bound L1 elements (i.e. L1 elements whose 5’UTR was <1 Kb from a distal – MLL2 peak losing H3K4me3 in double-KO cells) were identified using an in-house script. The percentage of each L1 subfamily with respect to all L1 elements were calculated for either the MLL2-bound L1 elements or all L1 elements.

## Data availability

All the genomic datasets generated in this project have been deposited into GEO. They will be publicly available upon peer-reviewed publication of this work.

## Supplementary figures

**Figure S1.**
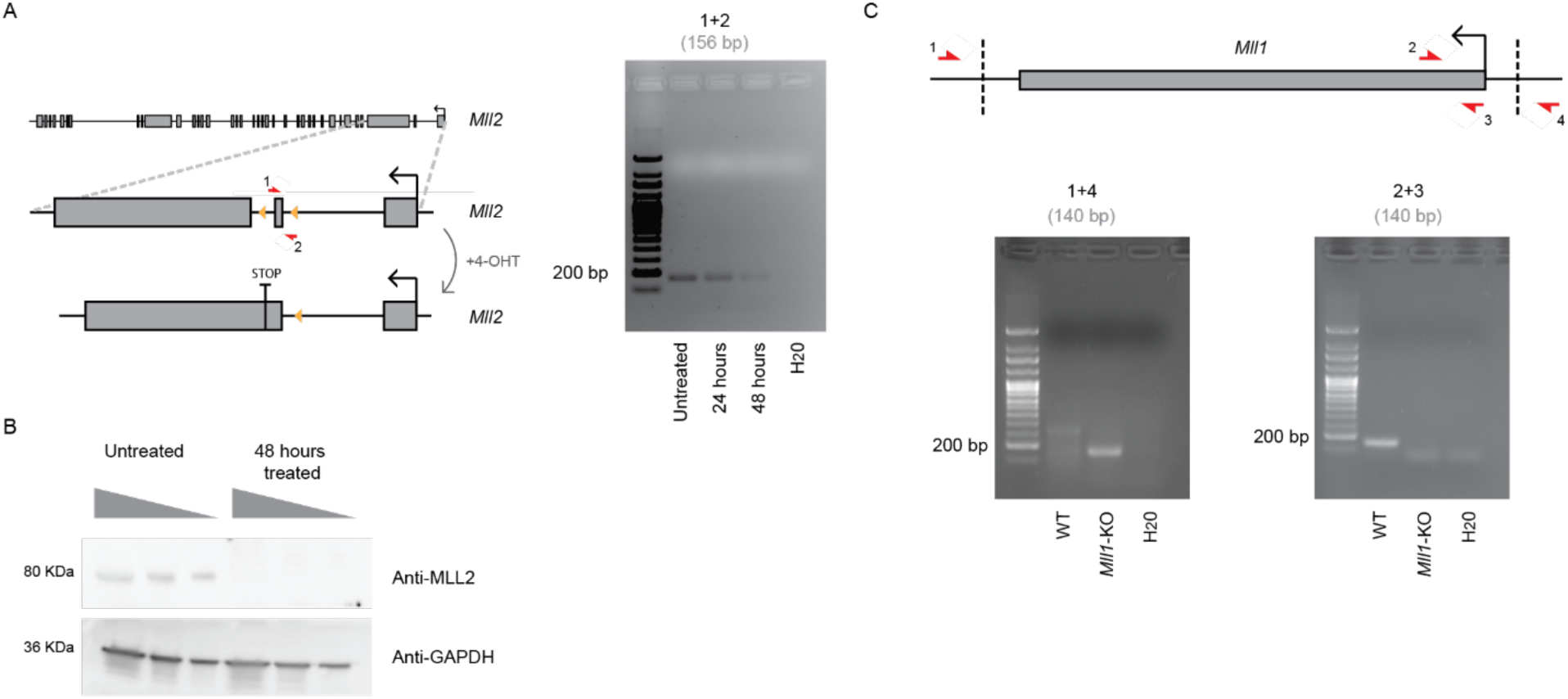
Generation and characterization of Mll1-KO, Mll2-KO and double-KO mESC. (A) Schematic representation of the strategy employed to generate inducible Mll2-KO mESC (left). Yellow triangles indicate the loxP sites flanking exon 2 of Mll2 in the conditional KO mESC line, which gets deleted upon addition of 4-OHT. Red arrows represent the primers used to assess the efficiency of exon deletion upon 4-OHT treatment by PCR (right). (B) Western blot showing MLL2 protein levels in Mll2-conditional KO ESC that were either untreated or treated with 4-OHT for 48 hours. GAPDH was used as a loading control. (C) Schematic representation of the strategy used to generate constitutive Mll1-KO mESC via CRISPR-Cas9 (upper panel). Dotted lines indicate gRNA target sites, and red arrows indicate the primers used to genotype the Mll1-KO mESC lines by PCR (lower panel).

**Figure S2.**
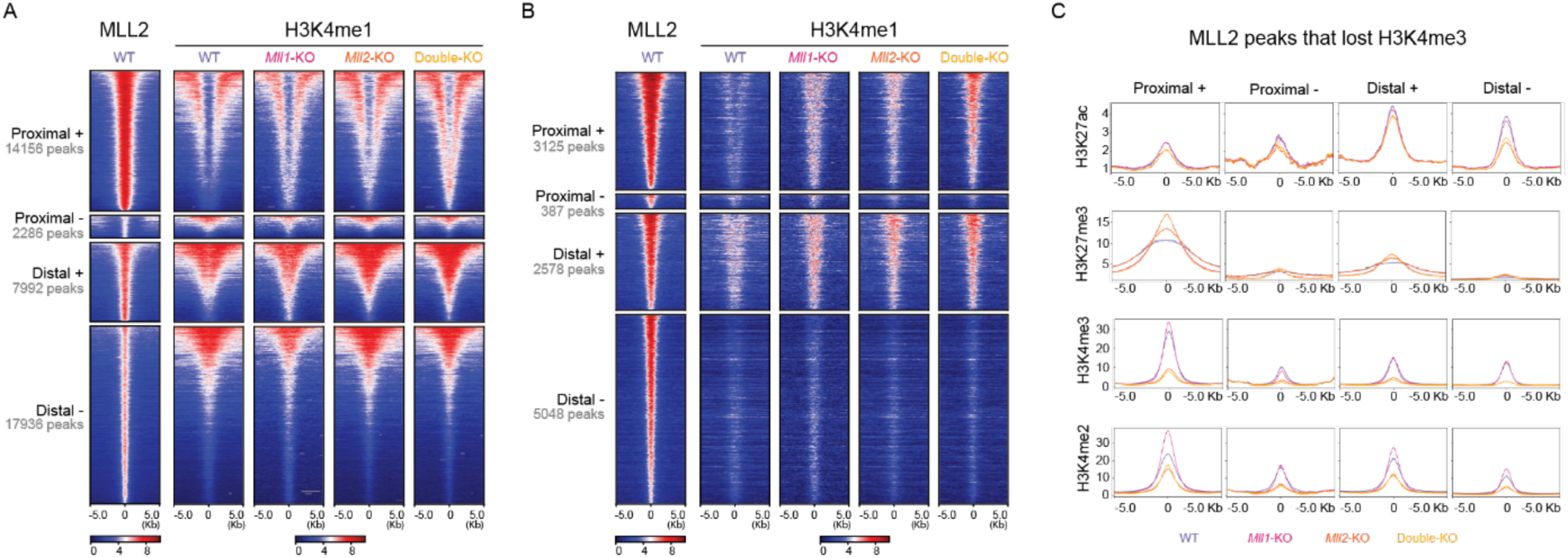
Epigenetic characterization of MLL2-bound regions in mESC. (A) Heatmap plots showing the ChIP-seq signals for MLL2 and H3K4me1 in WT, Mll1-KO, Mll2-KO and double-KO mESC around all the MLL2 peaks identified in WT mESC. MLL2 peaks were categorized in proximal +/- and distal +/- as described in Fig. 1B. Within each category, the MLL2 peaks were ranked according to H3K27ac signals in WT ESC (Fig. 1B). (B) Heatmap plots showing the ChIP-seq signals for MLL2 and H3K4me1 in WT, Mll1-KO, Mll2-KO and double-KO mESC around those MLL2 peaks that lost H3K4me3 in the double-KO mESC. The MLL2 peaks were categorized into proximal +/- and distal +/- as described in Fig. 1B. Within each category, the MLL2 peaks were ranked according to H3K4me3 signals in WT ESC (Fig. 1C). (C) Average ChIP-seq signal profiles for the indicated histone marks in WT, Mll1-KO, Mll2-KO and double-KO mESC around the MLL2 peaks that lost H3K4me3 in double-KO mESC. The MLL2 peaks were categorized into proximal +/- and distal +/- as described in Fig. 1B.

**Figure S3.**
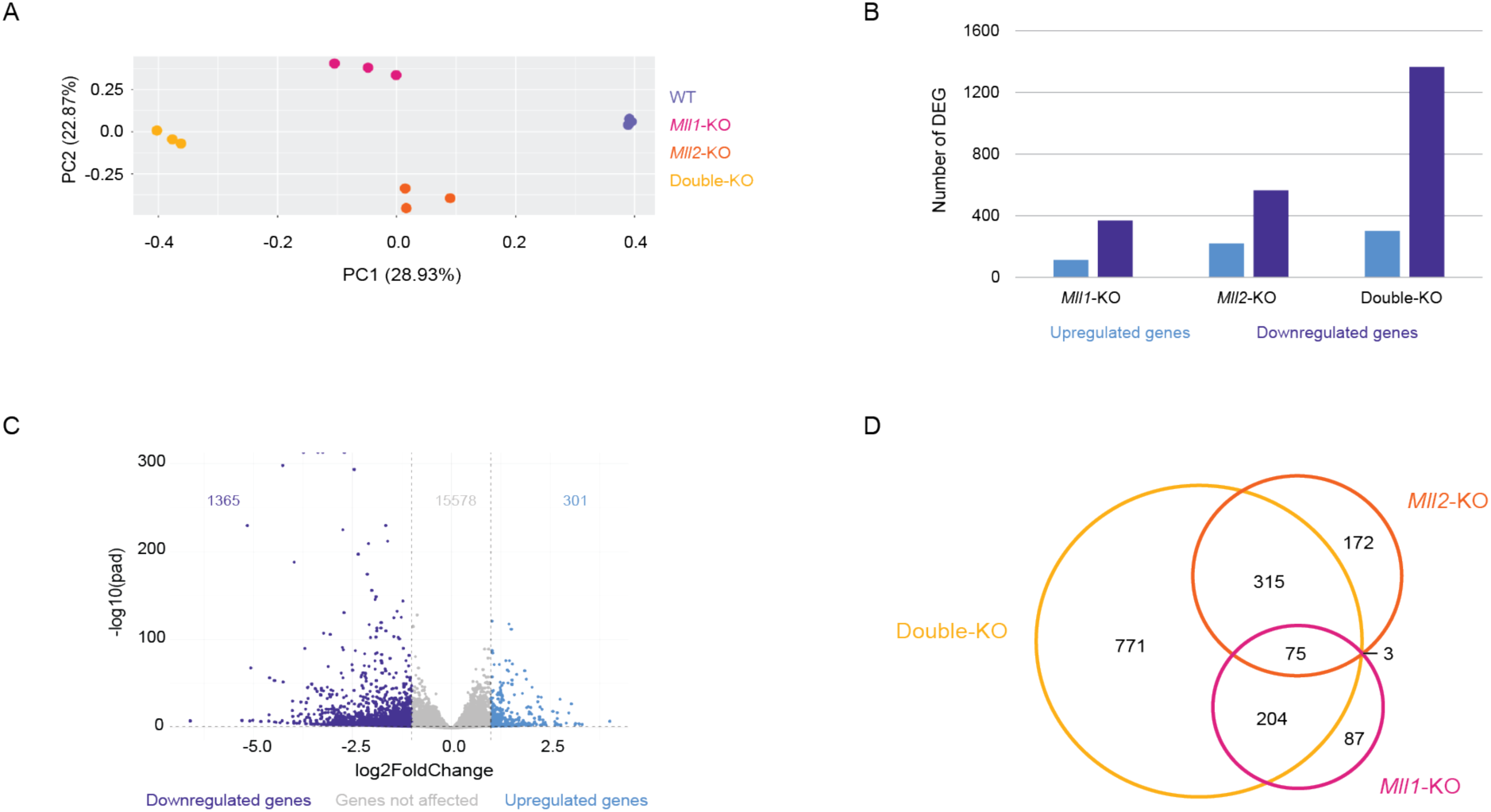
Gene expression changes following MLL2 loss in mESC. (A) Principal component analysis (PCA) of the RNA-seq datasets generated in WT, Mll1-KO, Mll2-KO and double-KO mESC. Three biological replicates were generated for each mESC line. (B) Barplot showing the number of differentially expressed genes in Mll1-KO, Mll2-KO and double-KO mESC compared to WT mESC. (C) Volcano plots illustrating global transcriptional changes in double-KO vs WT mESC. Each dot represents one gene and the genes that are either downregulated or upregulated in double-KO mESC are shown in purple and blue, respectively. The dashed horizontal line indicates a p-value = 0.05 and the dashed vertical lines indicate log2 fold-change = | 1 |. (D) Venn diagram showing the overlap among the genes downregulated in Mll1-KO, Mll2-KO and double-KO mESC compared to WT mESC.

**Figure S4.**
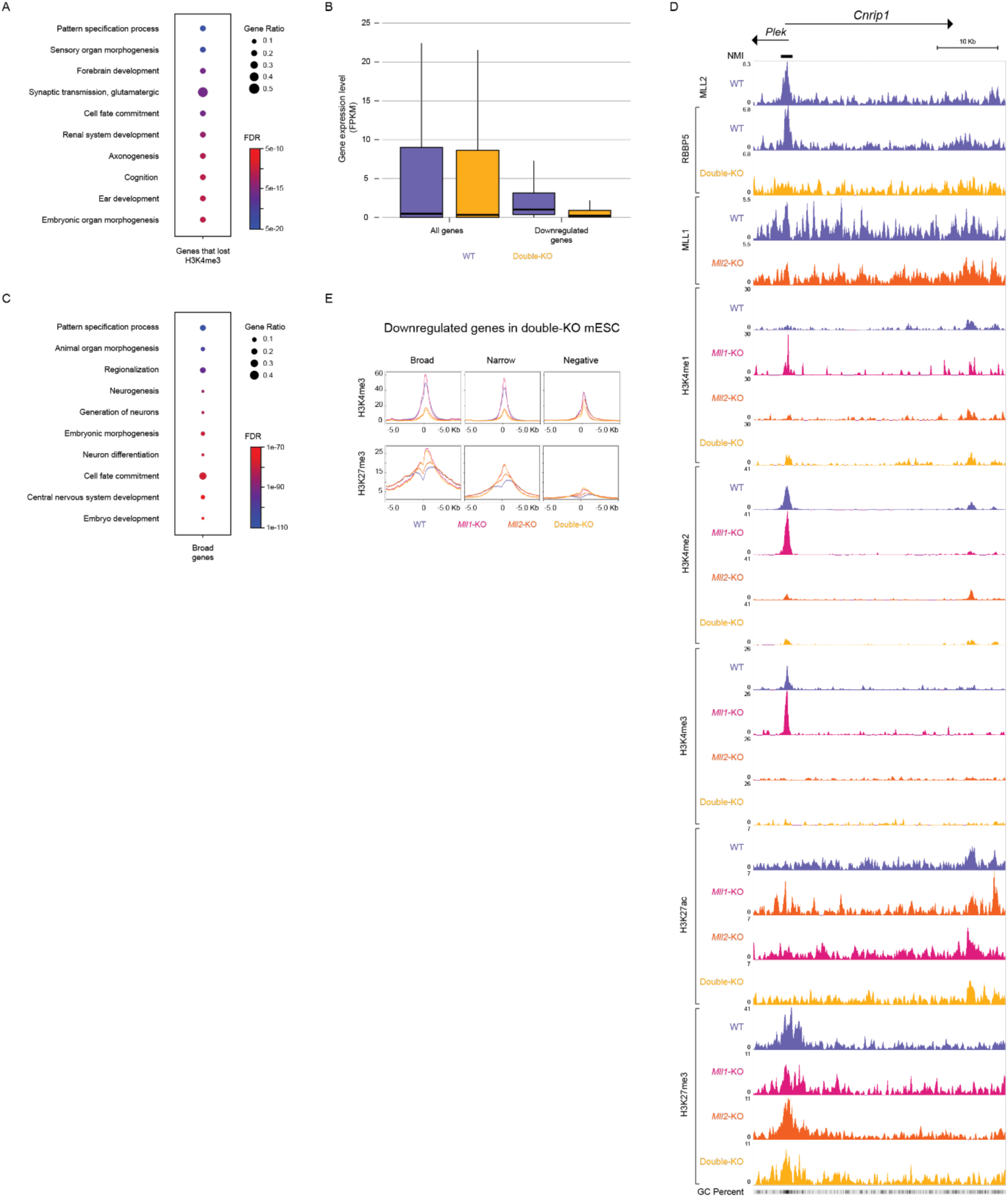
Epigenomic and transcriptional characterization of Mll1-KO, Mll2-KO and Double-KO mESC. (A) GO enrichment analysis for the “biological process” category was performed using WebGestalt for genes that lost H3K4me3 in double-KO mESC. Representative examples from the top 20 most enriched terms are shown. (B) Boxplot comparing gene expression levels (as measured by RNA-seq; FPKM) in WT and double-KO mESC for either all mouse genes or for genes downregulated in the double-KO mESC (vs WT mESC). (C) GO enrichment analysis for the “biological process” category was performed for genes associated with broad Polycomb domains using WebGestalt. Representative examples from the top 20 most enriched terms are shown. (D) Genome browser snapshot showing the ChIP-seq signals for the indicated TrxG proteins (i.e. MLL2, MLL1 and RBBP5) and histone marks in WT, Mll1-KO, Mll2-KO and double-KO mESC around Cnrip1, a gene downregulated in double-KO mESC. (E) Average H3K4me3 and H3K27me3 signal profiles in WT, Mll1-KO, Mll2-KO and double-KO mESC around the TSS of genes downregulated in double-KO ESC. The downregulated genes were divided into three categories based on the length of their associated Polycomb domains: broad (≥6 Kb), narrow (0< length <6 Kb) and negative (0 Kb).

**Figure S5.**
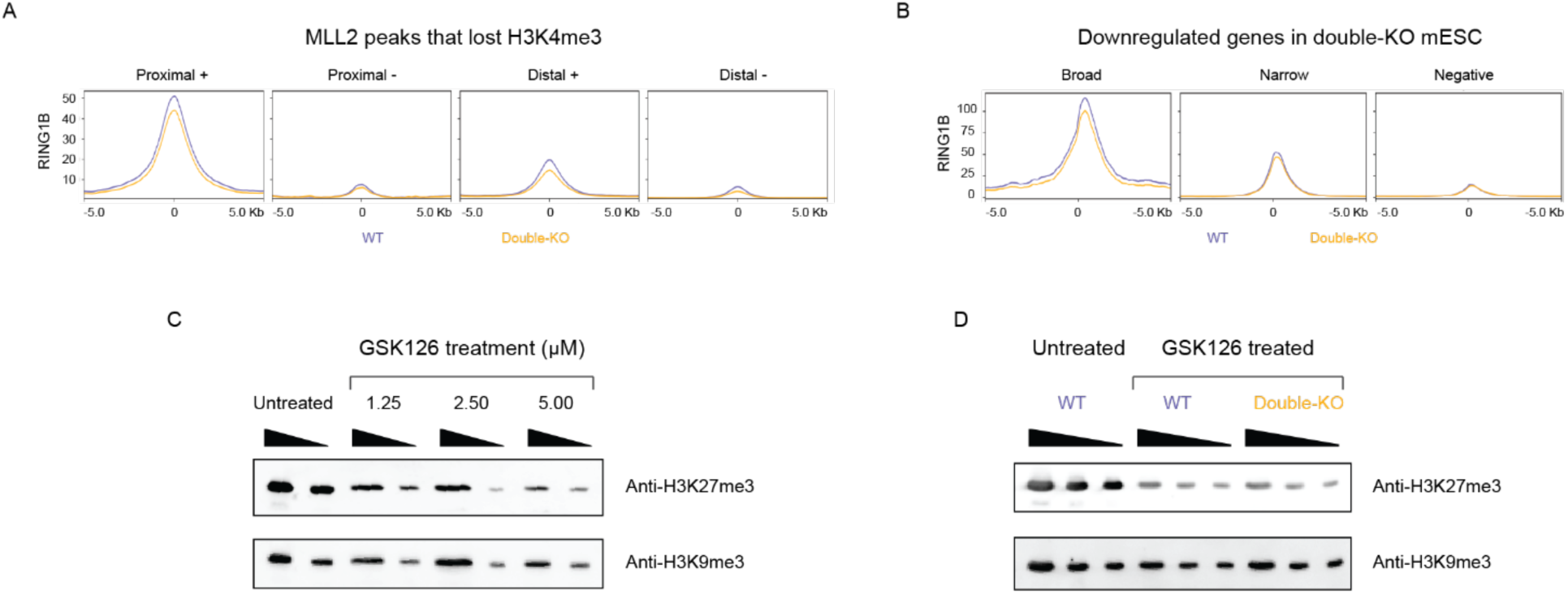
Evaluation of PRC1 and PRC2 repressive function in double-KO mESC. (A) Average RING1B ChIP-seq profiles in WT and double-KO mESC around the MLL2 peaks that lost H3K4me3 in double-KO mESC. The MLL2 peaks were categorized into proximal +/- and distal +/- as described in Fig. 1B. (B) Average RING1B ChIP-seq profiles in WT and double-KO mESC around the TSS of genes downregulated in double-KO ESC. The downregulated genes were divided into three categories based on the length of their associated Polycomb domains: broad (≥6 Kb), narrow (0< length <6 Kb) and negative (0 Kb). (C) Representative western blot from GSK126 dose optimization experiments performed in WT mESC. Cells were treated with 1.25, 2.5 and 5 µM GSK126 for 72 hours. Histone acidic extracts were prepared and analyzed for H3K27me3 levels. H3K9me3 was used as a loading control. (D) Western blot analysis of H3K27me3 and H3K9me3 levels in WT and double-KO mESC after 72-hour treatment with 5 µM GSK126. H3KSme3 was used as a loading control.

**Figure S6.**
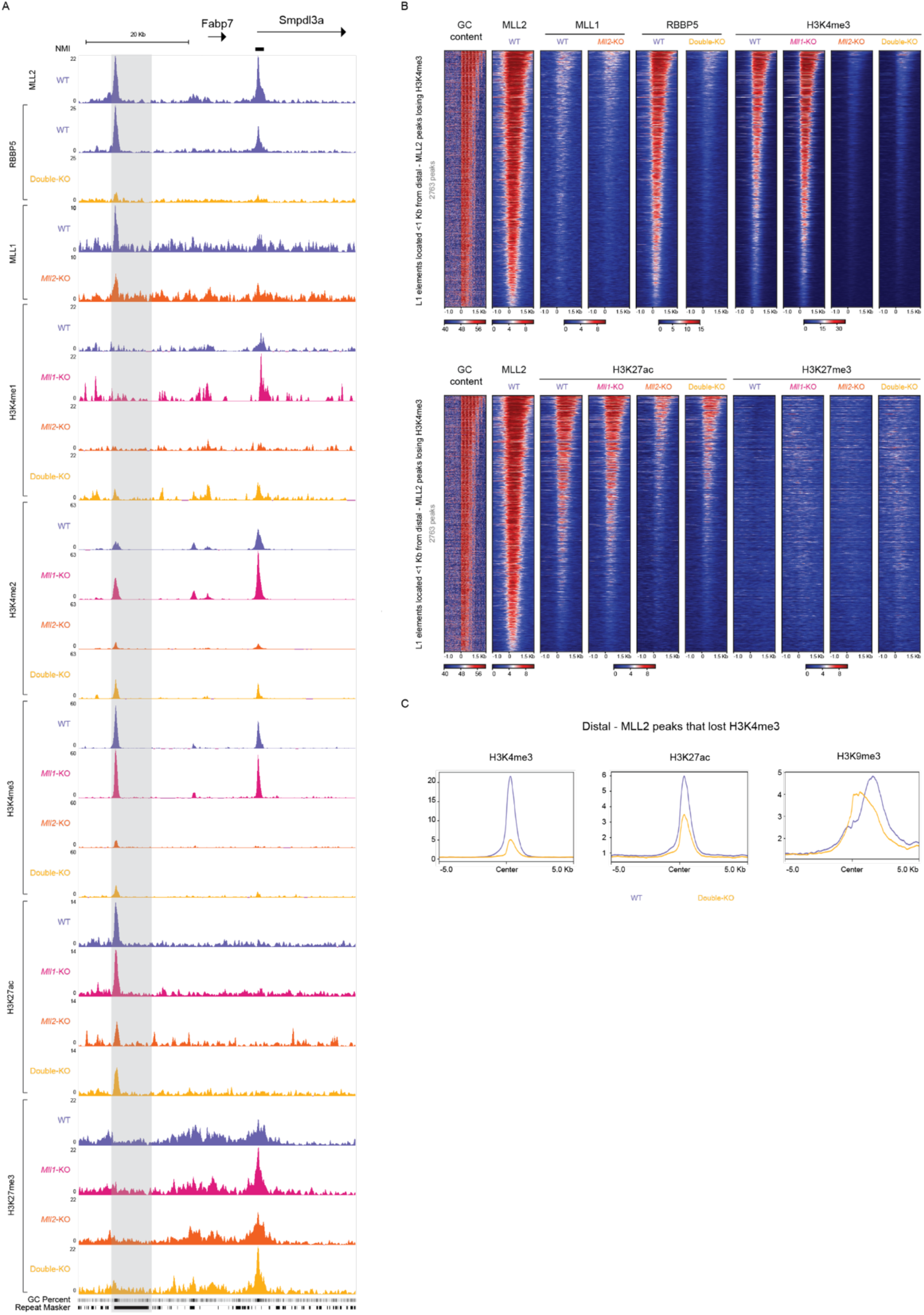
The 5’UTRs of MLL2-bound L1 elements are marked by active chromatin. (A) Genome browser snapshot showing the ChIP-seq signals for the indicated TrxG proteins (i.e. MLL2, MLL1 and RBBP5) and histone marks in WT, Mll1-KO, Mll2-KO and double-KO mESC around Smpdl3a, a gene downregulated in double-KO mESC. A nearby MLL2-bound L1 element is highlighted in grey. (B) Heatmap plots showing the GC percent and the ChIP-seq signals for the indicated TrxG proteins and histone marks in WT, Mll1-KO, Mll2-KO and double-KO mESC around the 5’UTR of MLL2-bound L1 elements (i.e. L1 elements located <1 Kb from distal - MLL2 peaks losing H3K4me3 in double KO mESC (n=2763)). L1s are sorted by the H3K4me3 signals in WT mESC. (C) Average ChIP-seq profiles for H3K4me3, H3K27ac and H3K9me3 around the 5’UTR of L1 elements located <1 Kb from distal - MLL2 peaks losing H3K4me3 in double KO mESC (n=2763).

**Figure S7.**
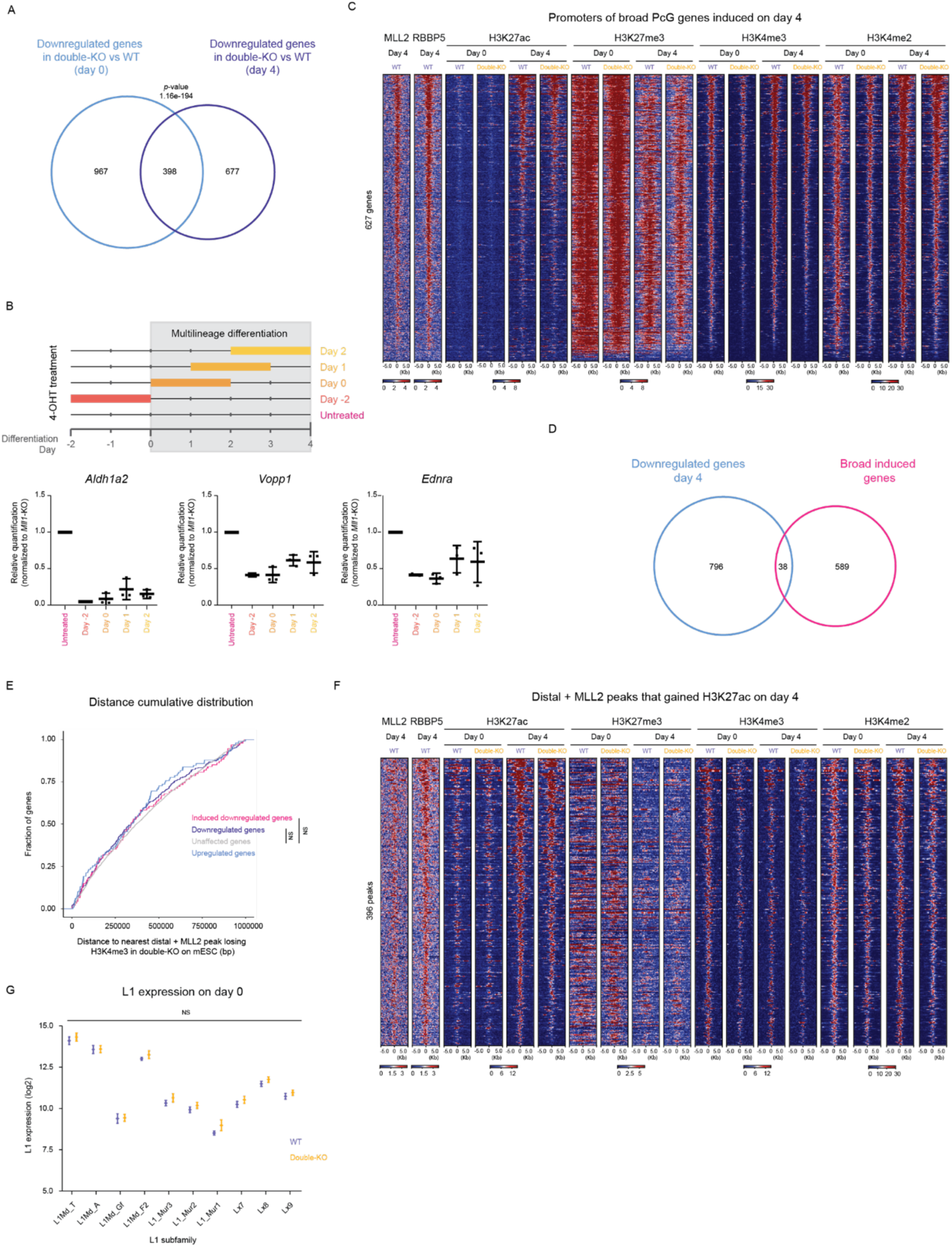
MLL2 is required for proper gene activation upon ESC differentiation. (A) Overlap between downregulated genes in double-KO cells in comparison to WT cells at either day 0 or day 4 of differentiation. The p-value was calculated using a hypergeometric test. (B) Time course analysis of 4-OHT treatment during multilineage differentiation. Upper panel: Mll1-KO cells were differentiation following a 4-day protocol. The 48 hours treatment with 800 nM 4-OHT to generate double-KO cells was initiated either 48 hours before the beginning of the differentiation (day -2), at the start of the differentiation (day 0), 24 or 48 hours after the differentiation was initiated (day 1 and day 2, respectively). Lower panel: expression levels of Aldh1a2, Vopp1 and Ednra in untreated and treated cells were assessed by RT-qPCR. Three biological replicates were collected for each condition, except for the day -2 condition (n=2). Expression values were normalized to untreated cells (i.e. Mll1-KO cells) using two housekeeping genes, Hprt and Eef1a1, as loading controls. (C) ChIP-seq heatmap plots around the TSS of genes with broad PcG-domains that got induced upon differentiation of WT ESC (day 4 vs day 0). ChIP-seq signals for the indicated TrxG proteins and histone marks are shown in WT and double-KO cells at day 0 and day 4. The TSS are sorted by the H3K4me3 signals in WT day 4 cells. (D) Overlap between downregulated genes in double-KO cells at day 4 (double-KO day 4 vs WT day 4; “Downregulated genes day 4”) and genes with broad PcG-domains that got induced upon differentiation of WT cells (WT day 4 vs WT day 0; “Broad induced genes”). (E) Cumulative frequency distribution plot for the distances from the TSS of unaffected, upregulated, downregulated and induced downregulated genes in day 4 double-KO cells to the nearest distal + MLL2 peak losing H3K4me3 in double-KO mESC. The “induced downregulated genes” category includes those genes that are downregulated in day 4 double KO cells in comparison to day 4 WT cells and that, in addition, are induced upon differentiation of WT cells (i.e. upregulated in WT day 4 vs WT day 0). NS: not significant. (F) ChIP-seq heatmap plots around distal + MLL2 peaks identified in WT mESC that gained H3K27ac in day 4 WT cells. ChIP-seq signals for the indicated TrxG proteins and histone marks are shown in WT and double-KO cells at day 0 and day 4. The MLL2 peaks are sorted by H3K27ac signals in WT day 4 cells. (G) The expression of the indicated L1 subfamilies in WT and double-KO mESC (i.e. day 0) was measured by RNA-seq using TEtranscripts^85^. For each L1 subfamily, the average expression levels and the corresponding standard deviation are plotted. The p-values were calculated using DEseq2 from TEtranscripts. NS: not significant.

**Figure S8.**
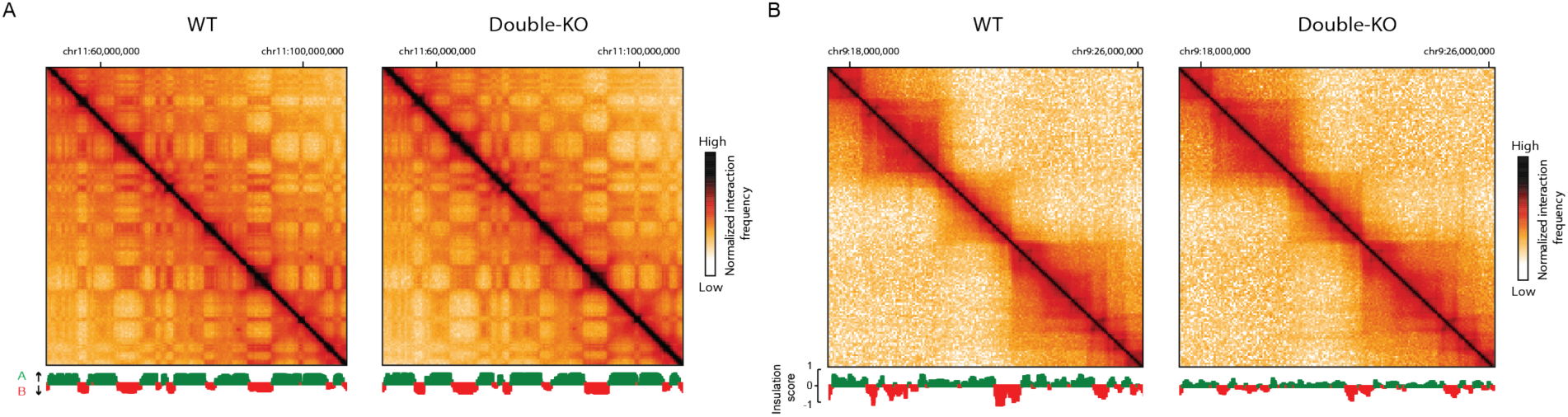
The loss of MLL2 does not significantly affect higher-order chromatin organization in mESC. (A) Micro-C normalized contact matrices for WT (left) and double-KO (right) mESC at 350 Kb resolution. Eigenvector values are shown at the bottom (positive/green values correspond to A compartments and negative/red values to B compartments). (B) Micro-C normalized contact matrices for WT (left) and double-KO (right) mESC at 50 Kb resolution. Insulation score values are shown at the bottom.

**Figure S9.**
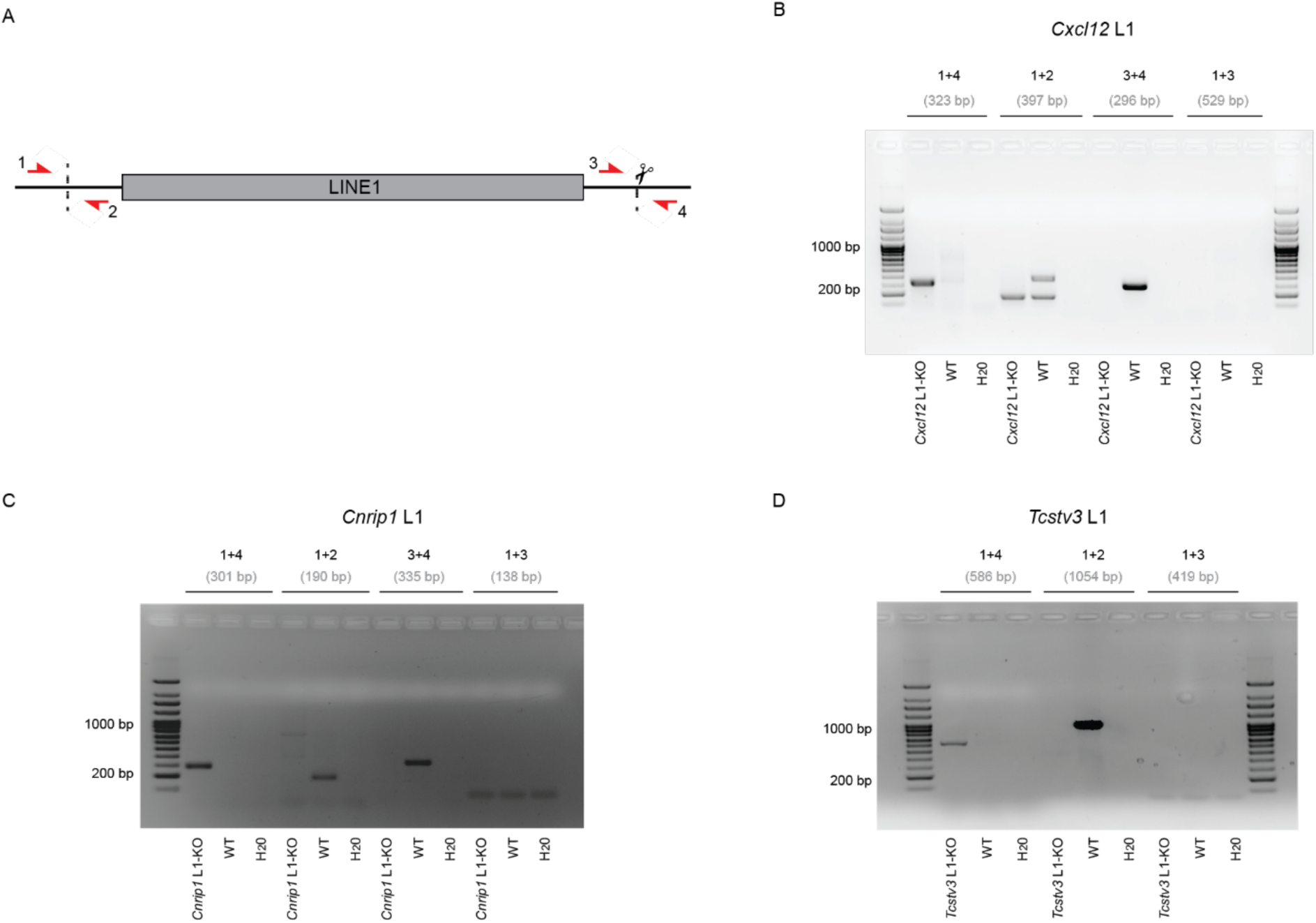
Generation and characterization of mESC lines with deletions of selected MLL2-bound L1 elements. (A) Schematic representation of the PCR-based strategy used to genotype the mESC lines with deletions of the selected MLL2-bound L1 elements. The dashed lines indicate the regions targeted by the gRNAs and the red arrows represent the primers used for PCR-based genotyping. (B-D) Representative PCR-based genotyping results of WT mESC and mESC lines with homozygous deletions of the MLL2-bound L1 elements associated with (B) Cxcl12, (C) Cnrip1 and (D) Tcstv3. The expected product sizes and the corresponding primer combinations are indicated above each lane. For the L1 element associated with Tcstv3, genotyping using primers 1+3 was not possible due to the repetitive nature of the genomic region flanking that side of the L1 element.

## Notes

### Competing Interest Statement

The authors have declared no competing interest.

